# Protein scaffold-based multimerization of soluble ACE2 efficiently blocks SARS-CoV-2 infection *in vitro* and *in vivo*

**DOI:** 10.1101/2021.01.04.425128

**Authors:** Alisan Kayabolen, Ugur Akcan, Dogancan Ozturan, Hivda Ulbegi-Polat, Gizem Nur Sahin, Nareg Pinarbasi Degirmenci, Canan Bayraktar, Gizem Soyler, Ehsan Sarayloo, Elif Nurtop, Berna Ozer, Gulen Guney-Esken, Tayfun Barlas, Ismail Selim Yildirim, Ozlem Dogan, Sercin Karahuseyinoglu, Nathan A. Lack, Mehmet Kaya, Cem Albayrak, Fusun Can, Ihsan Solaroglu, Tugba Bagci-Onder

## Abstract

Soluble ACE2 (sACE2) decoy receptors are promising agents to inhibit SARS-CoV-2, as their efficiency is less likely to be affected by common escape mutations in viral proteins. However, their success may be limited by their relatively poor potency. To address this challenge, we developed a large decoy library of sACE2 fusion proteins, generated with several protease inhibitors or multimerization tags. Among these decoys, multimeric sACE2 consisting of SunTag or MoonTag systems, which were originally utilized for signal amplification or gene activation systems, were extremely effective in neutralizing SARS-CoV-2 in pseudoviral systems and in clinical isolates. These novel sACE2 fusion proteins exhibited greater than 100-fold SARS-CoV-2 neutralization efficiency, compared to monomeric sACE2. SunTag or MoonTag in combination with a more potent version of sACE2, which has multiple point mutations for greater binding (v1), achieved near complete neutralization at a sub-nanomolar range, comparable with clinical monoclonal antibodies. Pseudoviruses bearing mutant versions of Spike, alpha, beta, gamma or delta variants, were also neutralized efficiently with SunTag or MoonTag fused sACE2(v1). Finally, therapeutic treatment of sACE2(v1)-MoonTag provided protection against SARS-CoV-2 infection in an *in vivo* mouse model. Overall, we suggest that the superior activity of the sACE2-SunTag or sACE2-MoonTag fusions is due to the greater occupancy of the multimeric sACE2 receptors on Spike protein as compared to monomeric sACE2. Therefore, these highly potent multimeric sACE2 decoy receptors may offer a promising treatment approach against SARS-CoV-2 infections.

**One Sentence Summary:** Multimerization of sACE2 markedly enhanced the neutralization of SARS-CoV-2 by blocking multiple viral spike proteins simultaneously.

## INTRODUCTION

Coronavirus disease 2019 (COVID-19) is a disease caused by severe acute respiratory syndrome coronavirus 2 (SARS-CoV-2) (*1*). It was firstly diagnosed in late 2019 in Wuhan, China (*2*) and has since spread over the globe, leading the WHO to declare a pandemic on March 11, 2020 (*3*). As of December 2021, there have been more than 270 million reported cases of COVID-19, which have caused more than 5.3 million reported deaths worldwide (*4*). Many vaccines have been developed in a remarkably short period of time and shown to be highly effective in phase III clinical trials (*5*–*7*) and real-life (*8*–*11*). Until now, 8 different vaccines have obtained emergency use listing (EUL) from WHO (*12*), and more than 8 billion doses of vaccines have been administered all over the world (*4*). While the vaccines are extremely effective in preventing severe illness and hospitalization rates, producing vaccines in sufficient amounts, and distributing them worldwide to stop the disease is still a challenging process. Indeed, there are countries with very low vaccination rates, either for economical, logistical, or individual reasons (*13*–*15*). As the new infections continue throughout world, emergence of novel variants of concern (VOC) such as the delta (*16*) or the omicron variants (*17*), is inevitable. Even single mutations found in new variants may increase the pathogenicity and binding potential of the virus and reduce the effects of vaccines (*18*–*20*). Therefore, effective drugs that specifically target SARS-CoV-2 and all its possible variants are still in unmet need to provide a reliable treatment.

For a long time, viral replication inhibitors such as Remdesivir or Favipiravir, which were initially developed for Ebola or influenza, respectively, were used as standard of care in the treatment of COVID-19 (*21*, *22*). However, recent clinical trials indicated little or no effect of these drugs especially for patients with severe symptoms (*23*, *24*). Recently, another viral replication inhibitor Molnupiravir developed originally for influenza treatment, was shown to have promising effects to reduce hospitalization and death rates in mild to moderate cases (*25*). Although there are numerous ongoing studies and clinical trials, drugs specifically targeting SARS-CoV-2 have not yet been identified or approved for clinical use.

The cell surface receptor responsible for SARS-CoV-2 infection is Angiotensin-converting enzyme 2 (ACE2) (*26*–*30*). Upon binding to ACE2, SARS-CoV-2 enters host cells via cleavage of its Spike glycoprotein by either membrane proteases such as TMPRSS2, or endosomal proteases such as cathepsin B/L (*29*). Camostat mesylate, which inhibits TMPRSS2, and E64d, which inhibits cathepsin B/L proteases, were found to be effective to neutralize SARS-CoV-2 (*29*). Accordingly, inhibitions of these key proteins were offered to block virus entry, and their efficacy is being tested in clinical trials (*31*). Similarly, natural or synthetic antibodies against ACE2 (*32*–*37*) or SARS-CoV-2 Spike (*38*), have been developed to prevent SARS-CoV-2 Spike binding to cellular ACE2.

However, SARS-CoV-2 was shown to acquire escape mutations under such evolutionary pressure, in line with viral adaptation mechanisms (*39*). Among the many variants that have emerged so far, some have been declared as VOCs due to their potential risks to public health. These variants were shown to have increased transmissibility or virulence, and resistance to vaccines or drug treatments because of escape mutations (*18*–*20*). Therefore, soluble ACE2 (sACE2) and its more potent forms have been proposed as promising alternatives to neutralizing antibodies (*40*–*42*), as they would prevent viral entry and further development of resistance. sACE2 was shown to inhibit SARS-CoV-2 infection in cell culture (*43*), *in vitro* human organoids (*43*), and recently a COVID-19 patient with severe symptoms (*44*). However, sACE2 is not the natural receptor of SARS-CoV-2 spike protein; therefore, high concentrations of sACE2 are required to achieve sufficient levels of neutralization.

In this study, we aimed to increase neutralization efficiency by generating a large number of novel sACE2 fusion proteins, using alternative protein domains such as protease inhibitors or multimerization tags. Our focus was particularly on multimerization tags, protein scaffold-based SunTag and MoonTag systems, which were originally developed for signal amplification (*45*, *46*). Indeed, these revealed great efficiency when fused to sACE2. As these multimerization based sACE2 fusions prevented the binding of host ACE2 and spike protein of SARS-CoV-2, they also demonstrated successful neutralization against the VOCs. Together, our results suggest a potent therapeutic approach against COVID-19.

## RESULTS

### sACE2-fusion proteins were developed to perform a small scale pseudovirus neutralization screen

To achieve high neutralization efficiency, we adopted molecular engineering and developed several sACE2-fusion proteins (**Figure S1**, **Table S1**). These were engineered based on different strategies including co-targeting of viral particles and host factors, and protein multimerization around virion. These fusion proteins included either sACE2(WT), the wild type form of sACE2, or sACE2(v1) that bears H34A, T92Q, Q325P, A386L point mutations and binds SARS-CoV-2 Spike with much higher affinity(*40*). For co-targeting strategy, we fused the extracellular domains of PAI1/*SERPINE1* or A1AT/*SERPINA1*, as potential TMPRSS2 inhibitors (*47*), and the extracellular domain of cystatin *C/CST3*, which was shown to inhibit cathepsin B/L in the endosomal pathway (*48*, *49*), to the C terminus of sACE2(WT) or sACE2(v1) via a GS rich linker (**Figure 1A**). In parallel, we hypothesized that protein-scaffold based multimerization of sACE2 would facilitate virus neutralization as it allows simultaneous binding of multiple Spike proteins around the virus. To test this hypothesis, we first utilized a multimerization system called SunTag, which was originally developed for signal amplification in fluorescence imaging (*45*). This system consists of two tags; a small GCN4 domain, a transcription factor found in yeast, and a scFv domain that is specific to the GCN4 tag. When SunTag was originally discovered, up to 24xGCN4 tags were used to allow recruitment of 24 copies of GFP (*45*). However, as sACE2 is significantly larger than GFP, we instead used a 5xGCN4 tag that had previously been incorporated in a dCas9-based gene activation system to recruit 5 copies of TET1 (*50*), which have a similar size with sACE2. In our design, we separately incorporated the 5xGCN4 or scFv-GCN4 tags to sACE2 (**Figure 1A).** Using both components together, we expected that sACE2 would be multimerized with these tags and bind to the multiple Spike proteins simultaneously. Therefore, achieving higher neutralization levels would be possible by increasing the sACE2 presence on the SARS-CoV-2 virion.

**Figure 1.**
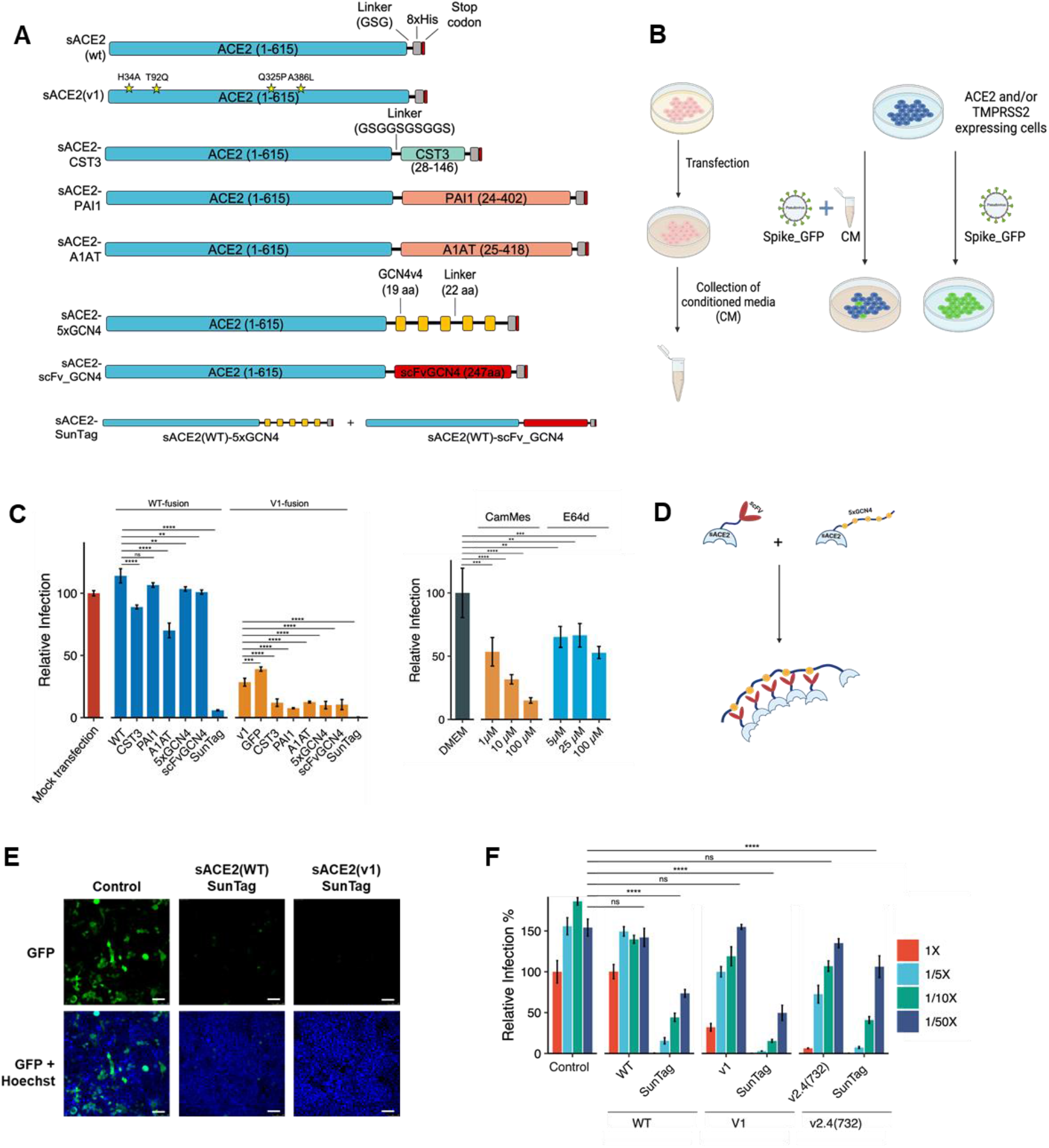
Pseudovirus neutralization screen with several versions of sACE2 fusions. **A)** Schematic representations of engineered sACE2-fusion constructs. **B)** Schematic model of the experimental approach conducted with conditioned media (CM) and pseudovirus bearing SARS-CoV-2 spike. (*Created with Biorender.com*) **C**) Relative infection rates of ACE2- and TMPRSS2-expressing HEK293T cells with pseudoviruses bearing SARS-CoV-2 Spike, in the presence of CM containing different sACE2 fusion constructs (*left*); or chemical inhibitors (*right*). CM from mock transfected cells were used as control (red bar), and infection rates were calculated as relative fluorescence to control wells. CamMes: Camostat Mesylate. **D)** Schematic representation of sACE2-SunTag. **E)** Representative images of pseudovirus neutralization via sACE2(WT)-SunTag and sACE2(v1)-SunTag. Pseudovirus-infected cells are shown in green. Cell nuclei stained with Hoechst 33342 dye are shown in blue. Scale bars = 100 μm. **F)** Relative infection with pseudovirus in the presence of different dilutions of CM containing different sACE2 constructs with or without SunTag. CM were diluted with DMEM up to 1/50X. CM from mock transfected cells were used as control, and infection rates were calculated as relative fluorescence to control wells. (ns: p > 0.05, *: p <= 0.05, **: p <= 0.01, ***: p <= 0.001, ****: p <= 0.0001, One way ANOVA)

### Pseudoviruses bearing SARS-CoV-2 Spike is a safe alternative to study virus neutralization *in vitro*

To test the neutralization efficiencies of designed constructs in a BSL-2 level laboratory, we generated lentiviral-based pseudotyped viruses containing a SARS-CoV-2 Spike glycoprotein and a GFP or luciferase transfer plasmids to measure cellular infection (**Figure S2A**). These Spike-bearing pseudoviruses were then used to infect HEK293T cells transiently expressing ACE2 and TMPRSS2. In parallel with the findings of Hoffmann et al. (*29*), cells co-expressing both ACE2 and TMPRSS2, were infected much more efficiently by GFP packaged Spike-bearing pseudoviruses, compared to cells expressing them individually (**Figure S2B and S2D**). As a control experiment, the infection efficiencies with VSV-G-bearing pseudoviruses were highly similar in all cell types (**Figure S2C and S2E**). Similar infection profile to GFP packaged Spike-bearing pseudoviruses was obtained with firefly luciferase (fLuc) packaged Spike-bearing pseudoviruses as well (**Figure S2F**). We also compared two different codon optimized Spike constructs when generating pseudoviruses, including the full-length version or 18-aa truncated version. The infection rate was nearly 6 times higher with the 18-aa truncated Spike (**Figure S2D**). Demonstrating the specificity of this model, pseudovirus infection was blocked with recombinant human ACE2 protein (**Figure S2G**). Encouraged by these results, ACE2 and TMPRSS2 co-expressing cells, and pseudoviruses bearing 18-aa truncated Spike were used for further experiments.

### Small scale pseudovirus screen revealed that SunTag mediated multimerization of sACE2 dramatically increased neutralization efficiency

Following the cloning of fusion proteins, we expressed them in HEK293T cells. As all fused proteins have extracellular domain of ACE2 in their N termini, they were secreted into the culture media. Assuming a similar expression for all constructs, we collected serum-free conditioned media (CM) and performed a small screen to test for pseudovirus neutralization (**Figure 1B**). Infection of cells with GFP encoding pseudoviruses were initially confirmed by microscopy (**Figure S3A**) and then quantified by a fluorescent plate reader (**Figure 1C**), while fLuc-encoding pseudoviruses were quantified by a luminescence assay (**Figure S3B**). From this initial screen, we observed that sACE2(WT) did not significantly impact pseudovirus neutralization. However, similar to findings of Chan et al. (*40*, *41*), sACE2(v1) including H34A, T92Q, Q325P, A386L mutations, could significantly inhibit the cell entry of pseudoviruses. The CST3 and A1AT fusions slightly increased the effects of both sACE2(WT) and sACE2(v1). As they were not effective individually (Figure S3C), we conclude that fusion with sACE2 might enhance their activity by bringing them closer to their target enzymes. However, the most striking results were obtained with the SunTag system. Crude CMs from both sACE2(WT)-SunTag and sACE2(v1)-SunTag almost completely abolished pseudovirus infection (Figure 1C-1E). While the individual components, sACE2-5xGCN4 and sACE2-scFvGCN4, had similar activity to the untagged sACE2, the efficient pseudovirus neutralization occurred through multimerization of sACE2 (**Figure 1C-E**).

As a comparison with recent studies, we included small molecule inhibitors of TMPRSS2 and Cathepsin B/L (Camostat mesylate and E64d) (*29*) in our pseudotyped virus neutralization assay. These inhibitors were effective in the micromolar concentration range as expected. However, even at the highest concentrations (100 μM), they were not as potent as sACE2(WT)-SunTag or sACE2(v1)-SunTag CM in inhibiting virus entry, attesting to their superior efficacy (**Figure 1C and S3B**).

We observed that both sACE2(WT)-SunTag and sACE2(v1)-SunTag CMs blocked more than 90% of pseudovirus infection. To compare the dosage dependency, we next performed a limiting dilution assay (**Figure 1F**). sACE2v2.4(732) was also added to the dilution assay, as it was shown to be highly potent against Spike, because of the point mutations and the protected natural dimerization domain (*40*, *41*). As expected, without dilution, sACE2v2.4(732) was more effective than the sACE2(WT) or sACE2(v1) and its effects were similar to WT-SunTag and v1-SunTag. However, even after 1/5x dilution, both WT-SunTag and v1-SunTag neutralized pseudoviruses significantly better than the others and were still effective after 1/50x dilution (**Figure 1F**). We also fused sACE2v2.4(732) with SunTag components to test whether we could enhance its neutralization efficiency. As expected, v2.4(732)-SunTag neutralized pseudoviruses better than its untagged counterpart (**Figure 1F**). However, it was not as potent as v1-SunTag, again attesting to the marked efficiency of v1-SunTag.

### Purified sACE2-SunTag decoys neutralized pseudovirus and SARS-CoV-2 isolates in sub-nanomolar doses

As a follow up to the experiments conducted with conditioned media, we generated recombinant proteins *in vitro* using 8xHis tagged WT, WT-SunTag, v1, and v1-SunTag ACE2 (**Figure 2A**) followed by fPLC purification. *In vitro* generated monomeric sACE2(WT), had comparable activity (IC_50_ = 358 ± 114 nM) to commercially available recombinant sACE2 (IC_50_ = 275 ± 78 nM) (**Figure S2G**). On the other hand, the sACE2(WT)-SunTag exhibited more than 250-fold neutralization capacity (IC_50_ = 1.35 ± 0.24 nM) compared to the monomeric sACE2(WT). Further, in line with our results with CM, the purified sACE2(v1)-SunTag was the most efficient system, with a sub-nanomolar median (IC_50_ = 0.30 ± 0.07 nM) inhibitory concentration (**Figure 2B**).

**Figure 2.**
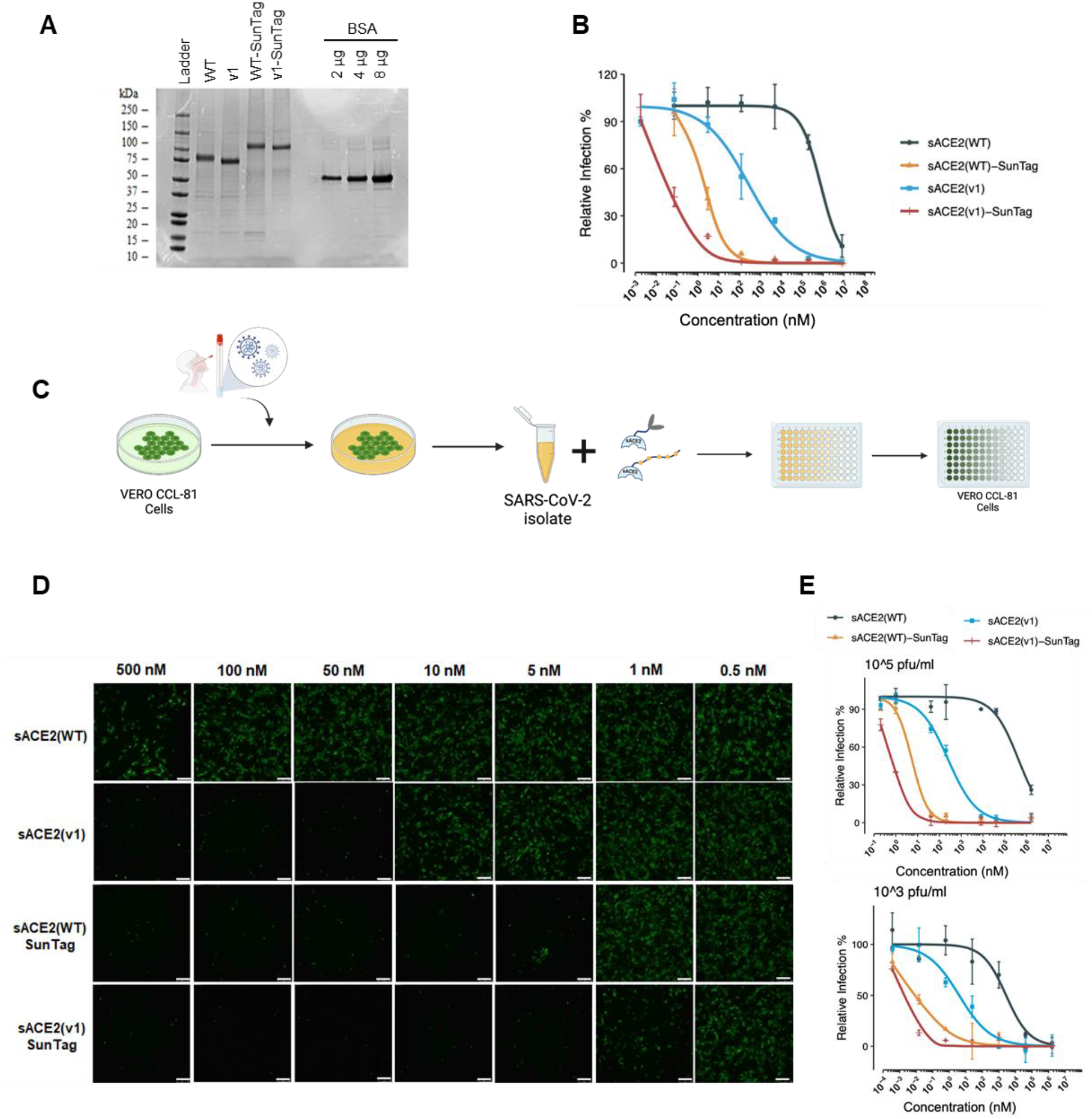
Neutralization of SARS-CoV-2 with sACE2 alone or sACE2-SunTag fusions. **A)** SDS-PAGE analysis of purified ACE2 (WT or v1) or their fusion proteins. Protein ladder and different amounts of bovine serum albumin (BSA) were used as controls. **B)** Relative infection rate of ACE2- and TMPRSS2-expressing HEK293T cells with pseudoviruses bearing SARS-CoV-2 Spike in the presence of purified sACE2 proteins. Purified proteins were serially diluted in culture media. Pseudovirus incubated with culture medium was used as control, and infection rates were calculated as relative fluorescence level of control. **C)** Experimental flow of SARS-CoV-2 isolation and neutralization effect of sACE2 fusions on Vero-CCL81 cells. (*Created with Biorender.com*) **D)** Microscopic images of Vero-CCL81 cells infected with SARS-CoV-2 isolate (10^5^ pfu/ml) upon 1h of incubation with different concentrations of purified sACE2-SunTag proteins. Cells were immunostained with anti-SARS-CoV-2 Spike antibody after 24h of infection (green). Scale bars =100 μm. **E)** Quantification of relative infection rate of Vero-CCL81 cells with 10^5^ pfu/ml or 10^3^ pfu/ml SARS-CoV-2 isolate upon treatment with purified sACE2 proteins. Infection was measured as fluorescence measured upon immunostaining with anti-SARS-CoV-2 Spike antibody.

To extend our findings from the pseudotyped virus model to a *bona fide* SARS-CoV-2 infection model, we carried out neutralization assays with clinical isolates of SARS-CoV-2 in a BSL-3 laboratory (**Figure 2C**). Accordingly, both sACE2(WT)-SunTag and sACE2(v1)-SunTag blocked SARS-CoV-2 entry to Vero-CCL81 cells much more efficiently than their untagged counterparts (**Figure 2D and S4**). sACE2(WT)-SunTag and sACE2(v1)-SunTag neutralized 10^5^ pfu/ml SARS-CoV-2 with IC_50_ values of 2.09 ± 0.45 nM and 0.84 ± 0.05 nM, respectively. Low dose infections with SARS-CoV-2 also responded to SunTag fusions at subnanomolar concentrations. While sACE2(WT)-SunTag neutralized 10^3^ pfu/ml with IC_50_ value of 0.15 ± 0.04 nM, sACE2(v1)-SunTag neutralized with IC_50_ = 0.06 ± 0.01 nM. (Figure 2E).

### MoonTag, another protein-scaffold based multimerization system, was markedly efficient in SARS-CoV-2 neutralization

Encouraged by our results with SunTag systems, we further tested the ability of a new multimerization system, MoonTag, which was developed to consist of two components: a linear peptide chain containing 15 amino-acid peptide repeats (gp41) and a nanobody (nb) specific to the gp41 peptide (*46*). We fused our sACE2 constructs with the MoonTag system components to generate multiple versions of MoonTag fusions. We fused both sACE2(WT) and sACE2(v1) with either 12xgp41 repeats or nb_gp41 (**Figure 3A**). Additionally, we fused 10xGCN4 and 24xGCN4 to sACE2(v1) to test different repeat numbers in the SunTag system. Using CM containing these fusions, we performed a pseudovirus neutralization assay to compare these novel multimerization systems with our previous SunTag system with 5xGCN4 component (**Figure 3B**). As expected, individual components of each system neutralized pseudovirus with a similar efficiency with the untagged version of sACE2(v1) (**Figure 3B and 1C**), while using components together neutralized pseudoviruses completely. We used 1/5x and 1/25x dilutions of CMs from multimerization systems as well, to compare their neutralization limits. 1/25x dilutions revealed that SunTag systems with 10xGCN4 and 24xGCN4 worked slightly better than SunTag with 5xGCN4 (**Figure 3B**). However, the most efficient neutralization was observed with the MoonTag system with 12xgp41.

**Figure 3.**
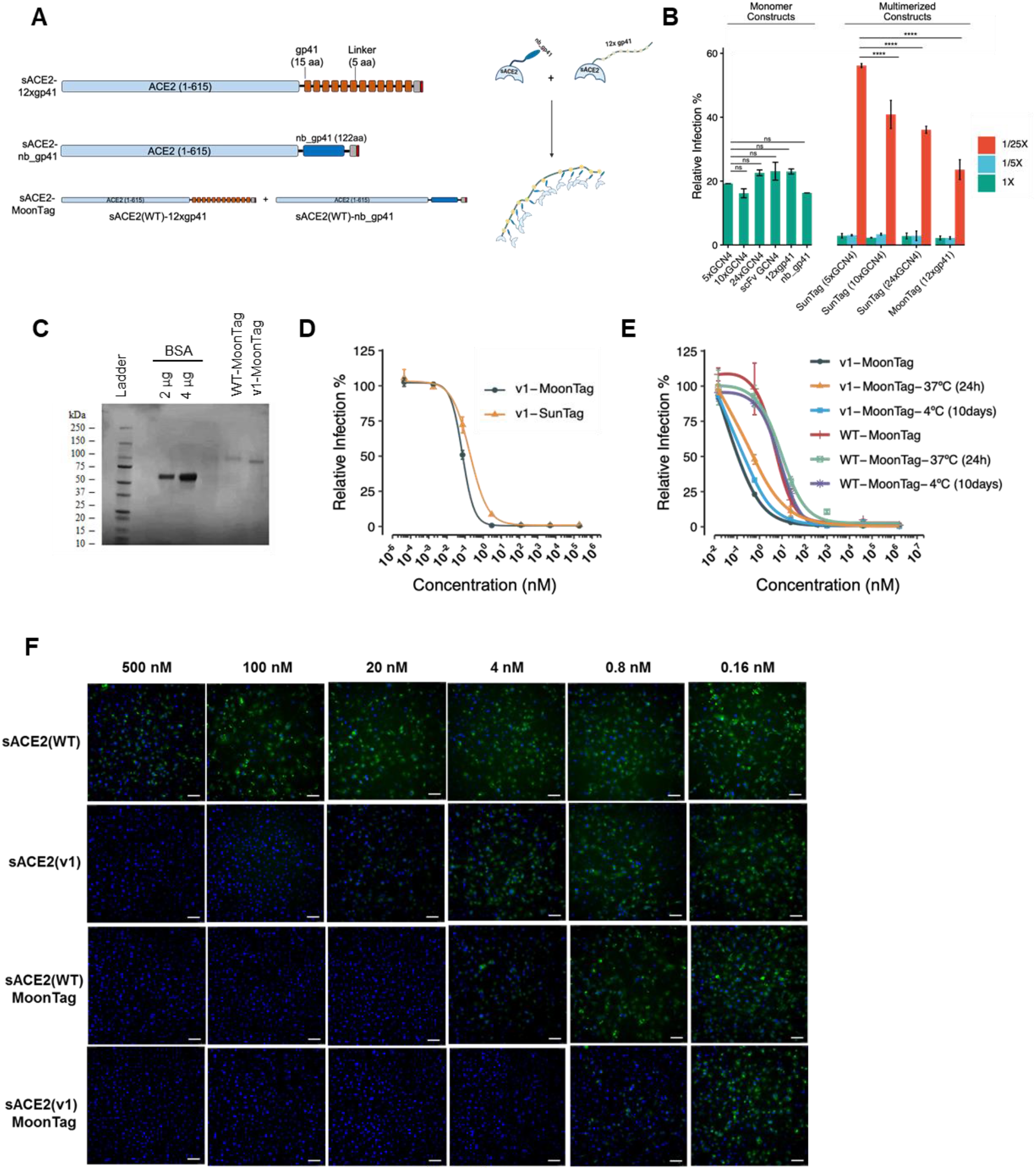
Pseudovirus and isolated SARS-CoV-2 neutralization with sACE2-Moon-Tag fusions. **A)** Schematic representation of sACE2 constructs with MoonTag. **B)** Relative infection of ACE2- and TMPRSS2-expressing HEK293T cells with pseudoviruses bearing SARS-CoV-2 Spike in the presence of CM containing sACE2 monomeric or multimerized fusion constructs. CM were diluted with DMEM up to 1/25X. **C)** SDS-PAGE analysis of purified MoonTag fusion proteins. Protein ladder and different amounts of bovine serum albumin (BSA) were used as controls. **D)** Comparison of SunTag and MoonTag-based sACE2-fusions for neutralization of SARS-CoV-2 Spike bearing pseudovirus. Relative infection rates were calculated as relative fluorescence intensity to virus-only conditions. **E)** Stability assay for sACE2(v1)-MoonTag. Pseudovirus neutralization assay was performed with purified v1-MoonTag proteins after incubation for 10 days at +4 °C or 24h at 37 °C. **F)** Microscopic images of Vero-E6 cells infected with SARS-CoV-2 isolate (10^5^ pfu/ml) upon incubation with different concentrations of purified sACE2 constructs for 1h. Cells were immunostained with anti-SARS-CoV-2 Spike antibody after 24h of infection. (Green: SARS-CoV-2 Spike; Blue: DAPI). Scale bars = 100 μm.

Encouraged by its significant neutralization ability in CM, we next produced pure sACE2-MoonTag proteins (**Figure 3C**). When compared with the SunTag system, the neutralization efficiency of the MoonTag was slightly better. sACE2(v1)-MoonTag proteins neutralized pseudoviruses at IC_50_ = 0.30 ± 0.02 nM, while sACE2(v1)-SunTag did so IC_50_ = 0.48 ± 0.07 nM (**Figure 3D**). As stability can be a major issue for protein-based drugs, we tested the impact of storage conditions on pseudovirus neutralization, keeping the proteins at either 4°C for 10 days or 37°C for 24h (Figure 3E). Although a slight increase was observed in their IC_50_’s, both WT-MoonTag and v1-MoonTag were still highly efficient to neutralize pseudoviruses (**Figure 3E**). Then, we tested the efficiency of the MoonTag system for neutralization of clinical SARS-CoV-2 isolates as well (**Figure 3F**). Similar to results of the pseudovirus experiments, sACE2(v1)-MoonTag neutralized SARS-CoV-2 isolates in sub-nanomolar range, and sACE2(WT)-MoonTag increased neutralization efficiency more than 100-fold when compared to sACE2(WT) monomer (**Figure 3F**). Together, these results suggest that sACE2-MoonTag fusions provide great efficacy in SARS-CoV-2 neutralization.

### SunTag and MoonTag-based multimerization of sACE2 significantly enhances binding to SARS-CoV-2 Spike

We designed a Spike binding assay to quantify the relative accumulation of sACE2 on the surface-expressed Spike protein (**Figure 4A**). After tagging sACE2 constructs with superfolder GFP (sfGFP), we collected CM for each fusion protein and then incubated it with Spike-expressing HEK293T cells. MoonTag components could not bind the Spike efficiently when they were incubated individually (**Figure 4B and 4C**). However, when the components were incubated together, both sACE2(WT) and sACE2(v1) bound to Spike proteins much more efficiently than their individual counterparts (**Figure 4B and 4C).** We also observed that v1-sfGFP-MoonTag bound to Spike very quickly, in a few minutes, upon incubation with cells (**Movie S1**). Similarly, the SunTag system highly enhanced the Spike binding when its components were applied together, compared to individual components or untagged sACE2 versions (**Figure S5A**). Immunostaining with anti-SARS-CoV-2 Spike antibody indicated that sfGFP-fused sACE2-SunTag specifically bound to Spike-expressing cells (**Figure S5B**). In line with our previous results, individual components of the SunTag system, sACE2-5xGCN4 and sACE2-scFvGCN4, did not demonstrate increased spike binding occupancy (**Figure S5A**). Moreover, when individual components of the MoonTag system were labeled with different fluorescent proteins, overlay images showed that their binding was colocalized (**Figure 4D**).

**Figure 4.**
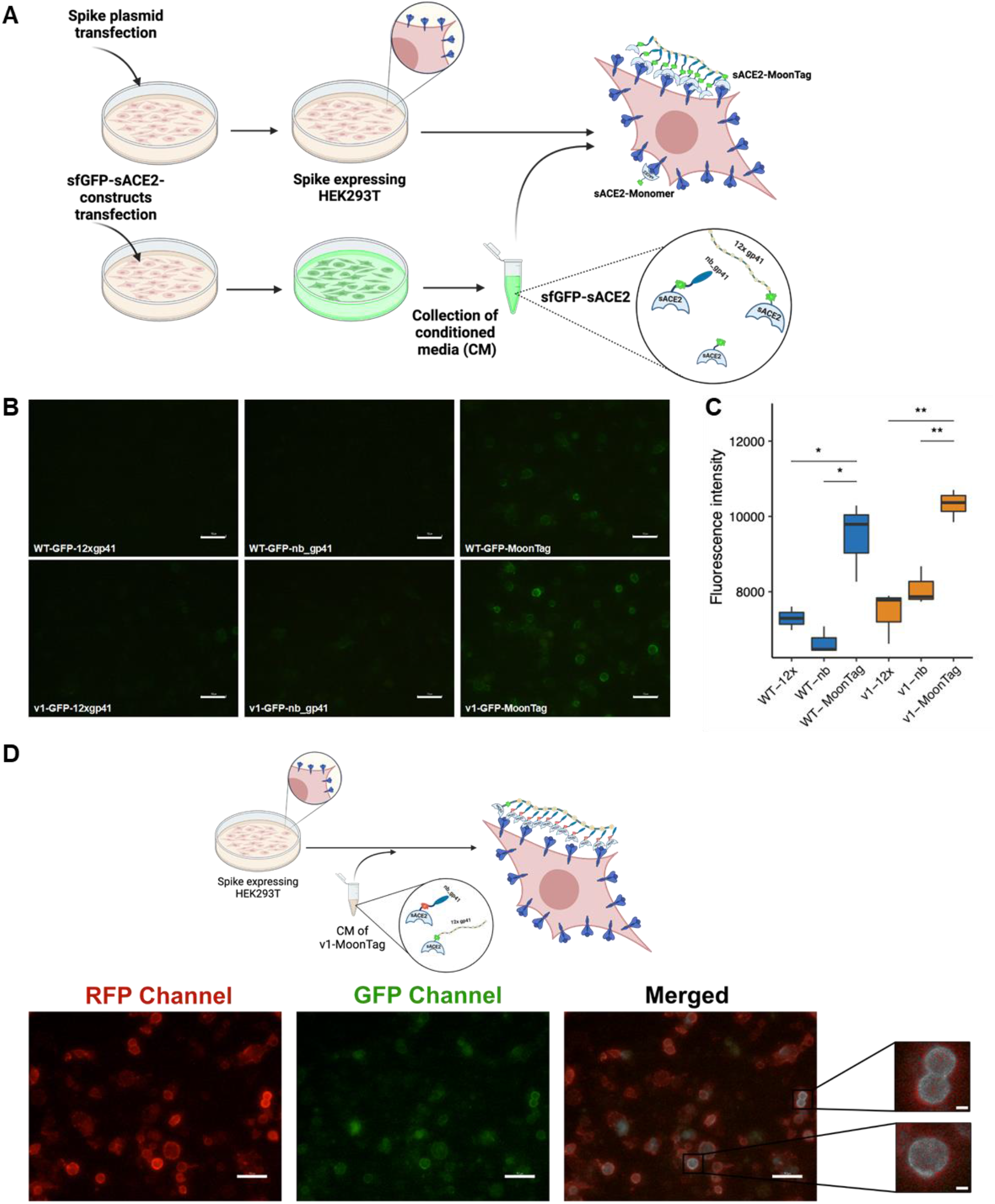
Spike binding assay with sACE2-MoonTag fusion proteins. **A)** Schematic representation of spike binding assay. (*Created with Biorender.com*) **B)** Microscopic images of Spike-expressing HEK293T cells upon incubation with CM collected from cells transfected with sfGFP-fused versions of sACE2(WT), sACE2(v1) with or without MoonTag components. Scale bars = 50 μm. **C)** Quantification of binding assay. Fluorescence intensities were calculated by analysis of 4 different images for each condition via ImageJ software. (ns: p > 0.05, *: p <= 0.05, **: p <= 0.01, ***: p <= 0.001, ****: p <= 0.0001, Student’s t-test.) **D)** Schematic representation and microscopic images of co-localization of MoonTag components. Spike-expressing HEK293T cells were incubated with CM of v1-MoonTag system containing mRFP-fused v1-nb-gp41 and sfGFP-fused v1-12x-gp41. Co-localized constructs were designated with white color in the merged image. Scale bars = 50 μm (10 μm in magnified insets). (*Created with Biorender.com*)

### MoonTag system exhibits significant neutralization efficiency against pseudo-variants of SARS-CoV-2

To test the efficiency of MoonTag system in neutralizing the VOCs of SARS-CoV-2, we produced pseudoviruses bearing SARS-CoV-2 Spike having 4 of the important mutations identified in alpha, beta, gamma, and delta variants (**Figure 5A**). Infection rates of all pseudo-variants were markedly higher than the pseudovirus bearing wild type Spike (**Figure 5B**). In neutralization assays, all pseudo-variants were neutralized by sACE2(v1)-MoonTag significantly (**Figure 5C**). Interestingly, IC_50_ for neutralizations of pseudo-variants, especially for the delta variant, was slightly lower than that of the wild type pseudovirus. In parallel with infection and neutralization data, Spike binding assay showed that binding affinity of sACE2(v1)-MoonTag to Spike variants is significantly stronger than binding to the wild type Spike (**Figure 5D and 5E**). This differential binding was more apparent with sACE2(WT)-MoonTag as the Spike variants can bind to wild type ACE2 more efficiently (**Figure S6).**

**Figure 5.**
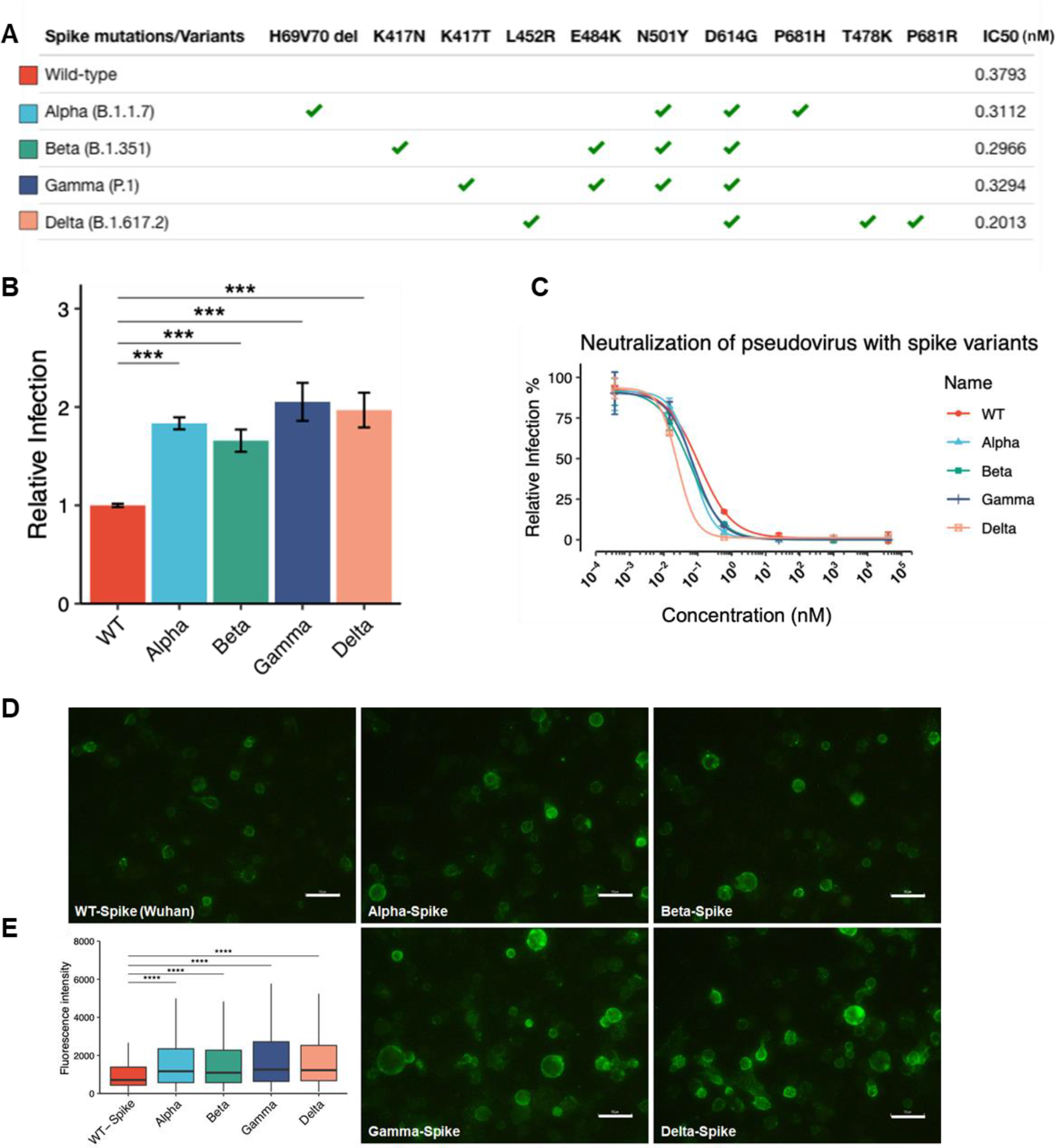
Neutralization of variants of concern (VOC) pseudoviruses and spike binding assay with sACE2(v1)-MoonTag construct. **A)** Representation of mutations engineered via site-directed mutagenesis for each VOC, and IC_50_ values for neutralization with purified v1-MoonTag proteins. **B)** Relative infection rate of ACE2- and TMPRSS2-expressing HEK293T cells with pseudoviruses bearing VOC Spike. Infection rates were normalized to fluorescence level of control cells infected with pseudoviruses bearing WT-Spike. **C)** Relative infection rate of ACE2- and TMPRSS2-expressing HEK293T cells with pseudoviruses bearing VOC Spike in the presence of purified sACE2(v1)-MoonTag proteins. Purified proteins were serially diluted in culture media. Culture medium was used as control, and infection rates were normalized to fluorescence level of control. **D)** Microscopic images of wild type or VOC Spike-expressing HEK293T cells upon incubation with CM collected from cells transfected with sfGFP-fused version of sACE2(v1)-MoonTag. Scale bars = 50 μm. **E)** Quantification of Spike binding assay in (D). 4 images from each condition were analyzed via ImageJ software. (ns: p > 0.05, *: p <= 0.05, **: p <= 0.01, ***: p <= 0.001, ****: p <= 0.0001, Student’s t-test.)

### sACE2(v1)-MoonTag is a potential inhibitor of SARS-CoV-2 *in vivo*

Based on our promising *in vitro* results, we tested the clinical potential of sACE2(v1)-MoonTag in a mouse COVID-19 model using K18-hACE2 transgenic mice expressing human ACE2. To mimic the natural disease, we infected the mice with SARS-CoV-2 virus (the alpha variant available in our labs) for 3 consecutive days, and beginning from 4h after the last infection, we administered purified sACE2(v1)-MoonTag or PBS as the control for 4 consecutive days (**Figure 6A**). After 7 days post infection (dpi), more than 10% average weight loss was observed in the control group, but there was no significant reduction in the average weight of the v1-MoonTag treated group (**Figure 6B**). At 8 dpi, 7 animals out of 8 in the control group died or were euthanized because of dramatic weight loss (>25%), while this number was only 1 out of 8 animals in the v1-MoonTag treated group (**Figure 6C**). At 9 dpi, while 3 animals in the v1-MoonTag treated group’s disease progressed, 4 animals (50%) were totally well and survived healthily until the end of the experiment (**Figure 6C**). Gross pathological analysis indicated that most of the lungs harvested from the control group have pneumonic and hyperemic regions, while the lungs harvested from healthy animals in the v1-MoonTag treated group were clear and had no indication of the infection (**Figure 6D**). Viral loads from the harvested lungs were also analyzed via qRT-PCR by using 2 primers targeting the viral nucleocapsid gene (**Figure 6E**). In parallel with the survival data, Cq values showed that 7 of the 8 animals in the control group had a high viral load (with Cq values < 20), while this number was 4 out of 8 animals in the v1-MoonTag treated group. The 4 healthy animals in the v1-MoonTag treated group had Cq values higher than 30. The histological examination of lung tissues revealed healing and restoration of normal tissue components in the v1-Moontag group (**Figure 6F, Figure S7**). The overall infiltration in the tissue decreased and normal tissue structure of alveolar spaces has been restored upon v1-MoonTag treatment. The interstitial and alveolar infiltrations declined to show only few inflammatory cells in the alveolar spaces. The exudate formed in the control group was not observed in the v1-MoonTag group. Increased number of type II pneumocytes and perivascular infiltration was noticed in the control group compared to v1-Moontag group demonstrating a mild perivascular and peribronchial infiltration. Together, these results show that protein scaffold multimerization based sACE2(v1)-MoonTag fusion reveals efficacy in the *in vivo* model of COVID-19.

**Figure 6.**
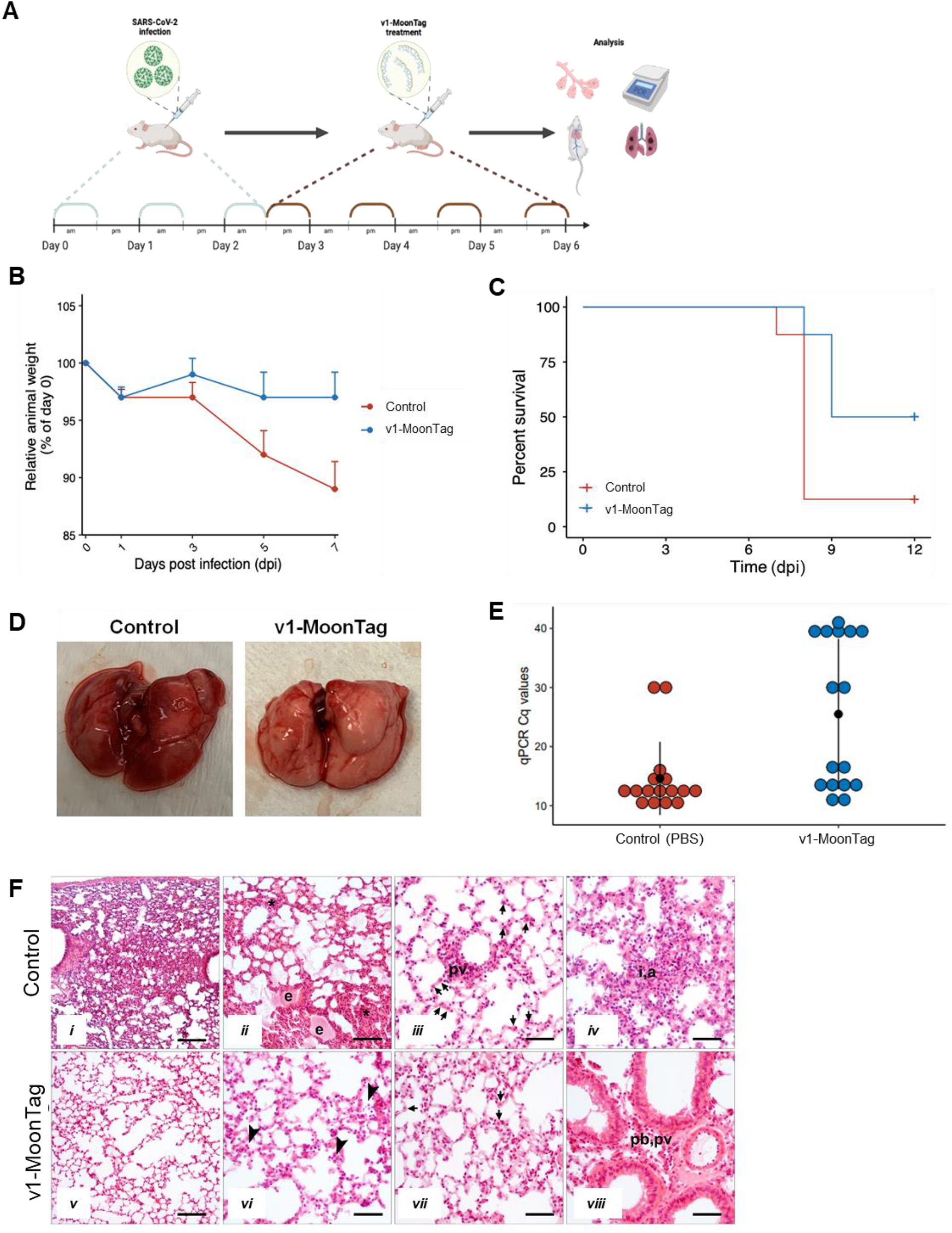
Neutralization effect of sACE2 constructs *in vivo*. **A)** Schematic representation of *in vivo* experiment to test efficacy of sACE2 constructs *in vivo*. Human ACE2 (hACE2)-expressing mice were infected with SARS-CoV-2 (Alpha) isolates for 3 days and treated with sACE2(v1)-MoonTag for 4 days starting from the 3rd day after infection (n=8/group). **B)** Average change in body weight by time in control or v1-MoonTag treated mice. **C)** Survival plot of animals in control and v1-MoonTag groups. **D)** Representative lung images dissected from control or v1-MoonTag treated mice. **E)** Viral loads in lungs from control or v1-MoonTag groups (n=8), measured via qRT-PCR from 2 different regions of SARS-CoV-2 Nucleocapsid gene. **F)** Representative images of hematoxylin-eosin stained lung tissues obtained from control (*i-iv*) and v1-MoonTag (*v-viii*) groups. The tissue architecture of v1-MoonTag group shows restoration of alveolar spaces with no exudate formation (*v*, *vi*); decreased amount of interstitial (*vi*), alveolar (*vii*), perivascular (pv, *viii*) and peribronchial (pb, *viii*) infiltration with a decline in number of inflammatory cells (arrowheads, *vi*); and re-establishment of a normal number of type-II pneumocytes (arrows, *vii*) compared to presence of exudate (e, *ii*); high amount of alveolar, interstitial (asterisk, *ii*; a,i, *iv*) and perivascular (pv, *iii*) infiltration along with an increased number of type II pneumocytes (arrows, *iii*) in the control group. Scale bars: *i*, *ii*, *v* = 70 μm; *vi*, *vii, viii* = 35 μm.

## DISCUSSION

COVID-19 pandemic is one of the most striking events of the last century. It caused millions of deaths and severely affected people’s lives in many ways including physical and psychological health, social relationships, and economic welfare. Vast amount of scientific research revealed crucial information about the disease and the virus, SARS-CoV-2, leading to the development of vaccines in a short time. Vaccines provide a high protection against the viral infection, minimize hospitalization and COVID-19 related deaths. However, difficulties in mass production and worldwide distribution, and vaccine hostility in many countries facilitates the emergence of novel SARS-CoV-2 variants. As the novel variants emerge, vaccine efficiencies are decreasing. Therefore, together with rapid and widespread vaccination programs, there is an urgent need for COVID-specific drugs, which can be used as curative to prevent disease progression. In this study, we provide novel sACE2-fusion proteins (**Figure 7**), that we developed using molecular engineering, as potent SARS-CoV-2 neutralizing agents that have potential as therapeutics against COVID-19.

**Figure 7.**
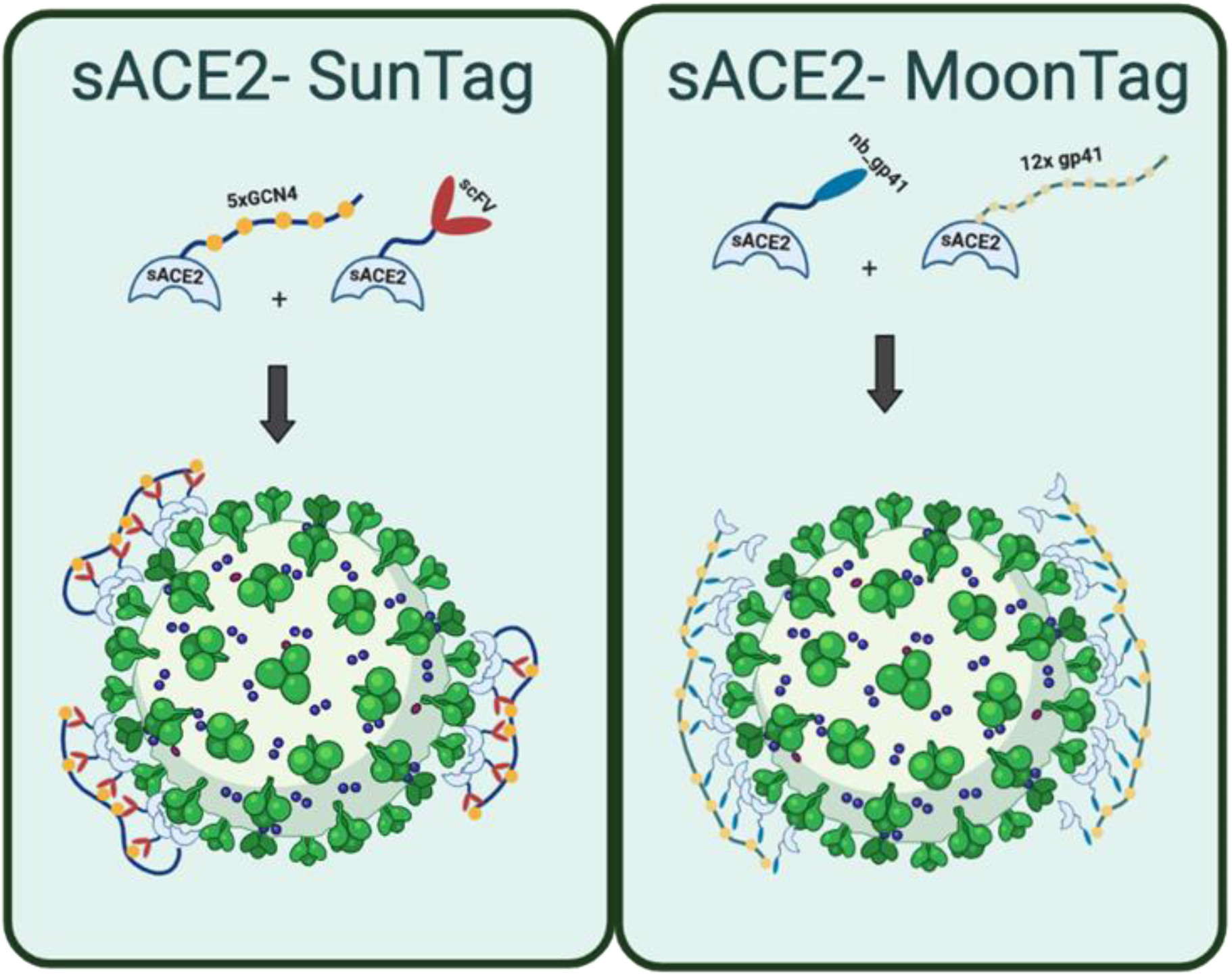
Graphical models of the Spike binding of multimerization-based sACE2-SunTag and sACE2-MoonTag fusion proteins.

There have been many treatment strategies against COVID-19, especially using antiviral agents such as Favipravir, Remdesivir or recently Molnupiravir (*23*–*25*). However, these agents do not specifically inhibit SARS-CoV-2 infection and exhibited less efficacy in recent clinical trials than originally thought (*23*, *24*, *51*). In parallel, several strategies based on Spike recognition with antibodies, peptides and small molecules have emerged as potential treatment options. However, it is known that mutations in novel variants, especially in the *Spike* gene, may hinder the recognition of Spike by antibodies (*52*, *53*); therefore, they may be less effective as escape mutations accumulate. Strategies to target ACE2 may be more feasible, since mutations that diminish ACE2 function would be detrimental to SARS-CoV-2 life cycle and therefore such mutations have not been identified. On the contrary, many common mutations found in variants were shown to increase the ACE2 binding affinity and increase the infection capacity of the virus (*54*–*57*). Therefore, using soluble ACE2 (sACE2) to block the natural ACE2-Spike binding is a promising approach as these soluble decoy receptors can potentially overcome common forms of SARS-CoV-2 resistance including the novel variants. However, the initial sACE2 decoy receptors have displayed relatively poor activity *in vitro* (*40*), and moderate effects in a recent clinical trial (*58*). To increase its potency, some research groups developed mutant forms of ACE2 with higher Spike affinity (*40*, *41*). Even though it is a successful strategy, it has the potential to cause development of novel escape mutations that can weaken the binding to mutant forms of ACE2 while alternatively strengthening the binding to wild type ACE2. Therefore, we designed a different strategy that increases the sACE2 potency without incorporating more mutations. We generated a small scale sACE2 decoy library with various fusion proteins that were categorized under two main approaches: co-targeting or multimerization (**Figure S1**, **S7, Table S1**).

In the co-targeting approach, we aimed to simultaneously block host ACE2-Spike interaction and cleavage of Spike by proteases. Both sACE2 decoy receptors and chemical inhibitors of host proteases such as TMPRSS2 or Cathepsin B/L were shown to be effective in preclinical studies (*29*). However, recent reports of clinical trials with these decoys or inhibitors revealed no significant effect on clinical outcome (*59*, *60*). To achieve better neutralization, we fused wild type or mutant (v1) sACE2 decoys with potential biological protease inhibitors, A1AT, PAI1, and CST3 (*47*–*49*). We hypothesized that the sACE2 component of these fusion proteins would block the Spike proteins on virions, and protease inhibitor components would prevent Spike cleavage even if some Spikes could bind the host ACE2. Our pseudovirus neutralization screen indicated that fusions with A1AT and CST3 increased the neutralization efficiency of sACE2. Indeed, A1AT was shown to be a potential inhibitor of SARS-CoV-2 infection recently (*61*). Therefore, its fusion to sACE2 might be a useful strategy to increase the neutralization potency.

In our second and more efficient strategy, we generated multimerization constructs for sACE2 decoys. Given that an average of 20-40 Spike trimers are found on a SARS-CoV-2 virion (*62*, *63*), most of these trimers should be blocked to neutralize the virion (*64*). We hypothesized that multimerization of sACE2 will facilitate neutralization as binding of one sACE2 molecule to any Spike protein would bring other sACE2 molecules closer to the other Spikes in the vicinity. We initially tested this hypothesis with a SunTag-mediated multimerization, and then followed up with a similar multimerization system, called MoonTag. Our results with both pseudovirus and clinical SARS-CoV-2 isolates clearly indicated that multimerization of sACE2 dramatically increased the binding affinity and neutralization efficiency, when compared to monomer sACE2. We investigated the applicability of these multimerization systems on both wild type sACE2, and its more potent mutant versions. SunTag or MoonTag-based multimerization of sACE2(v1) neutralized both pseudoviruses and SARS-CoV-2 isolates with sub-nanomolar IC_50_s, which are better than or comparable to several monoclonal antibodies developed recently (*32*–*34*).

This study is the first of its kind to examine the effects of varying different properties of multimerization system components on SARS-CoV-2 neutralization. We tested different numbers of peptide repeats in the tails, as 5x, 10x and 24xGCN4 in the SunTag system. Increasing the repeat numbers provided better neutralization at dilute concentrations, but this effect was limited after some point. This is probably because of lower expression of longer tails even with their higher multimerization capacity. However, all versions of tagged sACE2 decoys worked significantly better than the untagged sACE2 or individual monomer components. These results demonstrate that our approach of multimerization based sACE2 fusions are extremely efficient, in consistency with the recent studies indicating the success of dimerized (*40*) or trimerized (*42*) sACE2.

In addition to functional abilities, stability under different temperatures is an important issue for protein-based drugs. Therefore, we tested the neutralization capacities of our sACE2 fusion proteins upon storage at 4°C for 10 days, and at 37°C for 24h. Both sACE2(WT)-MoonTag and sACE2(v1)-MoonTag were highly stable in terms of their neutralization profile. Therefore, we suggest that these proteins could be amenable to current cold-chain logistics, be stored in conventional refrigerators for at least 10 days, and remain stable at physiological temperature for at least 1 day. Slight decrease in neutralization efficiency of sACE2(v1)-MoonTag upon storage at these conditions might be related with the point mutations it has. One of the four mutations found in sACE2(v1), T92Q, disrupts a consensus glycosylation motif formed by N90 and T92 residues which may lower overall stability (*40*). This may also be the reason why bands of v1 variants with or without multimerization tags were slightly weaker in SDS-PAGE compared to wild type sACE2. Therefore, using SunTag or MoonTag with other sACE2 variants (*40*–*42*), protecting the glycosylation motifs may give a better stability, and even better neutralization efficiency.

As multimerization systems offer to enhance neutralization efficiency by protecting the natural ACE2-Spike interaction, we suggest that this strategy will be similarly effective on any known or potential novel SARS-CoV-2 variants and provide experimental support with our engineered pseudoviruses bearing Spike variants, alpha, beta, gamma, and delta. Our sACE2-MoonTag fusions bound to and neutralized these variants, even with better efficacy compared to wild type Spike. Wild type sACE2 with MoonTag, especially, bound to variant Spikes much stronger than the wild type Spike. This is consistent with studies showing increased ACE2 affinity and infection capability of SARS-CoV-2 variants (*54*–*57*), and increased affinity and neutralization of variants with ACE2 decoys (*65*–*67*). These results clearly indicate that our multimerized sACE2 decoys are powerful tools against SARS-CoV-2 variants.

Our promising results *in vitro* encouraged us to test our multimerized sACE2 decoys *in vivo*, by using a COVID-19 model in mice. We selected sACE2(v1)-MoonTag for *in vivo* experiments since it was the most potent decoy *in vitro*. Interestingly, although many mutant, dimerized or trimerized ACE2 decoys were shown to have a high neutralization capacity *in vitro*, there are only a few *in vivo* studies showing efficacy of ACE2 decoys (*68*–*71*), or AAV-mediated ACE2 gene therapy (*72*). Although these studies show the potential protective effects of ACE2 decoys, most of them use a prophylactic approach and administer decoys/drugs prior to or together with viruses. A recent study testing the therapeutic potential of ACE2-Fc fusion, in which they administered proteins 12h after viral infection, obtained promising results in a hamster COVID-19 model (*70*). However, there are debates about fusions with Fc domains as it may cause severe immune response and increased viral entry via antibody-dependent enhancement (ADE) mechanism (*73*). To mimic the clinical conditions, we tested our decoys with a therapeutic approach, in which we administered sACE2(v1)-MoonTag after 3 days of SARS-CoV-2 virus (alpha variant) administration. We observed a significant protection of body weight and longer survival in the v1-MoonTag treated group compared to control. Based on clinical evaluation, qPCR, and survival results, we concluded that 4 of 8 mice in the v1-MoonTag treated groups were protected from disease progression at the end of the experiment. 3 of the 4 remaining mice had mild symptoms at 8 dpi but progressed at 9 dpi. To compare survival time between groups, we kept all mice alive as their clinical conditions were well, and harvested lungs after they got unhealthier. Therefore, viral loads in 4 completely healthy mice were expectedly low, while in others they were too high leading to a high error rate in the v1-MoonTag group. As our decoy treatment had ended at the end of 5 dpi, clinical outcomes might have been better if we continued with decoy treatment or increased the decoy dose. Changing the administration route from intraperitoneal to intranasal or intravenous injections might help achieve higher concentrations in lung tissue in future studies. All in all, complete protection of 4 of 8 mice supported with gross pathological and immunohistochemical images of healing in these mice indicate that sACE2(v1)-MoonTag is a promising drug candidate for COVID-19 treatment.

Although we obtained marked efficiency with multimerized sACE2 decoys both *in vitro* and *in vivo*, there are some limitations in our study. First, we could test our decoys on pseudo-variants, but not on clinical isolates of novel SARS-CoV-2 variants. Second, our *in vivo* study did not involve the monomer decoys or other potential multimer decoys to compare their effects in vivo. Challenges to produce large quantities of proteins, and to conduct animal experiments in ABSL-3 led us to minimize the experimental groups. Future studies will be performed to compare therapeutic efficacies of various sACE2 decoys with different doses and administration routes in vivo.

In this work, we demonstrate that incorporating multiple sACE2 via a scaffold-based multimerization system can generate a decoy receptor that can effectively prevent SARS-CoV-2 cellular infection (**Figure 7**). With sACE2(WT), this multimeric decoy receptor was more than 100-fold more active than the monomeric protein. Utilizing a mutant sACE2(v1), which has greater affinity for Spike, the neutralization efficiency with our multimeric system reached comparable levels to Spike monoclonal antibodies currently in clinical development. This proof-of-principle study utilized a SunTag or MoonTag approach to generate multimeric decoy receptors but any alternative peptide-scFv/nb couples could be incorporated with the ACE2. Overall, we suggest that highly potent multimeric sACE2 decoy receptors are promising agents, which can be used as a treatment for COVID-19 caused by any SARS-CoV-2 variants.

## MATERIALS AND METHODS

### Cloning of plasmids

5xGCN4 and scFvGCN4 multimerization tags were amplified from pPlatTET-gRNA2 plasmid, which was a gift from Izuho Hatada (Addgene #82559) (*50*). 12xgp41, 12xGCN4 and 24xGCN4 were amplified from 24xMoonTag-kif18b-STOP-24xSunTag-linker-24xPP7, which was a gift from Marvin Tanenbaum (Addgene plasmid # 128605) (*46*). Nb_gp41 was amplified from Nb-gp41-Halo (MoonTag-Nb-Halo) plasmid, which was a gift from Marvin Tanenbaum (Addgene #128603)(*46*). CST3 cDNA was amplified from CST3 (Homo sapiens) in pLenti6.3/V5-DEST (DNASU plasmid repository) (*74*). A1AT/ *SERPINA1* and PAI1/ *SERPINE1* cDNAs were amplified from SERPINA-bio-His plasmid (Addgene #52182) and SERPINE1-bio-his (Addgene #52077) which were gifts from Gavin Wright without signal sequence to prevent cleavage of fused proteins (*75*). All inserts were amplified with linker (GSGGSGSGGS) at the N-terminal, and with or without 8-his tags at the C-terminal, and cloned into BamHI and XbaI sites, instead of sfGFP either in pcDNA3-sACE2(WT)-sfGFP (Addgene plasmid, #145171), or pcDNA3-sACE2v1-sfGFP (Addgene plasmid, #145172) or pcDNA3-sACE2v2.4(732)-8h (Addgene plasmid, #154103), which are gifts from Eric Procko(*40*). sACE2(WT)-8his was generated via replacing sACE2v1 in the pcDNA3-sACE2v1-8his (Addgene plasmid, #149269) with sACE2(WT). sfGFP or mRFP fused sACE2 constructs were generated by cloning sfGFP or mRFP amplified with GS-rich linker, into BamHI site between sACE2 and 5xGCN4, scFvGCN4, 12xgp41, nb_gp41 domains in the sACE2(WT)-5xGCN4, sACE2(WT)-scFvGCN4, sACE2(v1)-5xGCN4, sACE2(v1)-scFvGCN4, sACE2(WT)-12xgp41, sACE2(WT)-nb_gp41, sACE2(v1)-12xgp41, sACE2(v1)-nb_gp41 plasmids. pCEP4-myc-ACE2 was a gift from Eric Procko (Addgene plasmid #141185). RRL.sin.cPPT.SFFV/TMPRSS2(variant 1).IRES-neo.WPRE (MT130) was a gift from Caroline Goujon (Addgene plasmid # 145843). pGBW-m4137383 was a gift from Ginkgo Bioworks (Addgene plasmid #149541). pcDNA3.1-SARS2-Spike was a gift from Fang Li (Addgene plasmid #145032) (*76*). All constructs were checked by diagnostic digestion and confirmed by Sanger sequencing of the insertion regions in plasmids. Details of all constructs generated for this study are outlined in Table S1.

### Site Directed Mutagenesis

Variants of concern (VOC) of SARS-CoV-2 were created with Q5® Site-Directed Mutagenesis Kit (New England BioLabs, Ipswich, MA, USA) according to manufacturer instructions. Spike-18aa truncated plasmid (Addgene plasmid # 149541) was used as template, and PCR reaction was performed with specific primers designed using NEBase Changer tool according to VOC mutations (Alpha: H69V70del, N501Y, D614G, P681H; Beta: K417N, E484K, N501Y, D614G; Gamma: K417T, E484K, N501Y, D614G; Delta: L452R, T478K, D614G, P681R). PCR products were treated with KLD (Kinase, Ligase & DpnI) enzyme mix to circularize them. Plasmids generated via SDM were transformed to *E. coli* (Stbl3) and isolated by using a mini prep kit (M&N, Germany).

### Cell culture

HEK293T cells were purchased from American Type Culture Collection (ATCC, USA), and cultured in DMEM (Gibco, USA), with 10% FBS (Gibco, USA) and 1% Pen/Strep (Gibco, USA) in a 37°C incubator with 5% CO_2_. Vero CCL-81 cells were also purchased from ATCC and cultured in DMEM with 5% FBS and 1% Pen/Strep in a 37°C incubator with 5% CO_2_. As HEK293T cells detach easily, tissue culture plates used for them were coated with 0.05% poly-L-lysine (PLL) for 30 min at 37°C and washed with PBS before seeding cells. For transfection experiments, HEK293T cells were seeded on 15-cm plates as 9×10^6^ cells/plate, 6 well plates as 1 × 10^6^ cells/plate, or 12 well plates as 0.4 × 10^6^ cells/plate. The next day, media were refreshed, and cells with approximately 70-90% confluency in each plate were transfected with required plasmid(s) for each assay using Fugene 6 transfection reagent (Promega, USA) or 1 mg/ml PEI (polyethyleneimine).

### Conditioned media preparation

Transfections were performed with Fugene 6 transfection reagent (Promega, USA) according to manufacturer’s instructions with minor modifications. Briefly, cells were seeded to reach %70-90 confluency at the day of transfection for Fugene and %90-100 confluency for PEI. 1:4 ratio of DNA/transfection reagent (Fugene or PEI) were mixed with DMEM without FBS or Pen/Strep (only DMEM) as the total volume will be 10 times of the transfection reagent and incubated for 5 min at room temperature. Total of 15 μg of fusion protein expression plasmid(s) for 15-cm plates, 2.5 μg for 6 well plates, and 1 μg for 12 well plates were prepared as separate mixture, added on the transfection reagent mixture, and incubated for 20-30 min at room temperature. Then, the mixture was added on cells dropwise. After 14-16h of transfection, media was removed, cells were washed with PBS, and only DMEM was added on cells. After 48 and 72h of transfection, conditioned media (CM) was collected from plates, centrifuged for 5 min at 1500 rpm, and supernatants were filtered with 0.45 μm syringe filters to remove cell debris. CM was freshly used or kept at +4°C for a maximum of 1-2 days.

### Pseudovirus production and infection

Lentiviral-based pseudoviruses bearing VSV-G or SARS-CoV-2 Spike (S) glycoproteins were produced as in previous studies with some modifications (*77*). Briefly, ~9×10^6^ of HEK293T cells were seeded on 15-cm plates, 1 day before transfection. Then, they were transfected with 7500 ng of lenti RRL_GFP reporter plasmid, 6750 ng of psPAX2 packaging plasmid (Addgene plasmid # 12260), and either 750 ng of Spike-18aa truncated (Addgene plasmid # 149541), or generated Spike variant plasmids (alpha, beta, gamma, or delta), or pcDNA3.1-SARS2-Spike (Addgene # 145032) or VSV-G plasmid (Addgene plasmid # 8454) as glycoprotein. After 14-16h of transfection, media was removed, and fresh media (DMEM with 10% FBS and 1% Pen/Strep) was added on cells. After 48h of transfection, viruses were collected, filtered through 0.45 μm syringe filters, and stored at +4 °C for short term usage (up to 3-4 days), or aliquoted and stored at −80 °C for long term storage. HEK293T cells were also transfected with ACE2 and TMPRSS2 expression plasmids (Addgene plasmid #141185 and #145843, respectively) individually or together, to allow infection by pseudoviruses. After 24h of transfection, cells were seeded on 96-well plates as 20k/well and infected with 50 μl of pseudoviruses (MOI ~ 1). The next day, viral media was replaced with fresh culture media.

### Neutralization assay for pseudovirus

50 μl of pseudoviruses bearing either wild type or variants of Spike were mixed with 50 μl of CM or purified proteins at different doses. For SunTag samples, 1 volume of CM from 5x-GCN4 was mixed with 3 volumes of CM from scFv-GCN4, and for MoonTag samples, 1 volume of CM from 12x-gp41 was mixed with 4 volumes of CM from nb-gp41 as the total volume is equal with CM from other samples. Mixtures were incubated for 30 min at 37°C temperature and used to infect ACE2- and TMPRSS2-expressing HEK293T cells. Infection rate by pseudoviruses were determined by fluorescence or luminescence reads in a microplate reader (BioTek’s Synergy H1, VT, USA), as GFP or fLuc-expressing reporter plasmids were packaged during pseudovirus production. Neutralization efficiency was calculated as relative fluorescence or luminescence to the CM collected from mock-transfected cells, or storage buffer for purified proteins.

### Protein purification

HEK293T cells were transfected with plasmids encoding required proteins as described in the CM collection procedure. CM were collected after 48h of transfection and filtered through 0.45 μm syringe filters and used for protein purification via ÄKTA pure 25 FPLC (GE Healthcare) equipped with 1-ml HisTrap affinity column (HisTrap HP, GE Healthcare) loaded with Ni (II) and previously equilibrated with PBS containing 5 mM imidazole (buffer A). Briefly, 45 ml of filtered CM from each sample was applied onto the column at the flow rate of 0.8 ml/min. After washing the column with 15 column volumes (CV) of buffer A at a flow rate of 2 ml/min, proteins were eluted with 15 CV of PBS containing 0.5M imidazole (buffer B) by a linear gradient (0 to 100% of buffer B at flow rate of 1 ml/min). The proteins were eluted in a peak centered at around 225 mM Imidazole concentration. The fractions containing the recombinant protein were pooled and concentrated through 15 mL 10-kDa MWCO Amicon-Ultra centrifugal filters (Millipore). Protein concentrations were measured by absorbance at 280 nm in nanodrop (Thermo Scientific). They were aliquoted as ~0.2 mg/ml and stored at −80 °C after snap freezing in liquid nitrogen.

For bulk purification of proteins to use *in vivo*, HEK293T cells were seeded on multiple 15-cm plates and transfected with plasmids encoding required proteins as described in the CM collection procedure. CM were collected after 48, 72, and 96h of transfection and filtered through 0.45 μm syringe filters and used for protein purification. CM were loaded into Amicon® Ultra Centrifugal filters (MWCO 10 or 30 kDa, Merck, Germany), and centrifuged at 3750 rpm for 30 minutes to concentrate CM and eliminate proteins that have low molecular weights. PBS was added into the filters continuously after each centrifugation until the phenol red was washed away completely. Concentrated samples were collected and loaded onto the homemade columns that were manually built-in syringes using Ni Sepharose™ High Performance beads placed on squeezed cotton wool. The unbound proteins were washed away with a wash buffer (PBS containing 7.5 mM imidazole, pH=8). Proteins were eluted from the columns using an elution buffer (PBS containing 250 mM imidazole, and protease inhibitors, pH=8). To eliminate imidazole, the elution buffer was exchanged with PBS using Amicon® Ultra Centrifugal filters, and proteins were concentrated by serial centrifugation. Protein concentrations were measured by absorbance at 280 nm in Nanodrop (Thermo Scientific). They were aliquoted as ~1-2 mg/ml and stored at −80 °C after snap freezing in liquid nitrogen.

### Isolation of authentic SARS-CoV-2

SARS-CoV-2 was isolated at Koç University Isbank Center for Infectious Diseases (KUISCID) Biosafety Level-3 (BSL-3) facility, from a nasopharyngeal sample of a confirmed COVID-19 patient admitted to the Koç University Hospital (approved by Koç University Ethics Committee with document number 2020.135.IRB1.025). Vero CCL-81 or E6 cells were cultured in Dulbecco’s Modified Eagle Medium (DMEM) supplemented with antibiotic/antimycotic and heat-inactivated 5% fetal bovine serum (FBS) were incubated with the nasopharyngeal sample for 4-5 days. After observation of cytopathic effect, the growth of SARS-CoV-2 was confirmed by qRT-PCR using primer and Taqman probe targeting SARS-CoV-2 nucleocapsid. To prepare virus stocks, Vero CCL-81 or E6 cells were infected, and supernatants were collected after cell debris removal by centrifugation at 48-hour post infection. Viral titer (TCID50) of the SARS-CoV-2 isolate was determined by Spearman-Karber method (*78*). To sequence the virus, viral RNA was extracted with Viral RNA isolation kit (QIAamp viral RNA), DNA libraries were prepared using the Illumina TruSeq stranded total RNA kit, and the viral RNA was sequenced using Illumina MiniSeq (GenBank: MT675956).

### SARS-CoV-2 neutralization assay

Purified soluble proteins were serially diluted in DMEM containing 5% FBS and antibiotic/antimycotic to generate required final concentrations. Equal volume of diluted proteins and 10^3^ plaque forming unit (PFU)/ml (MOI = 0.002) or 10^5^ PFU (MOI = 0.2) of SARS-CoV-2/ MT675956 were incubated for 1 hour at 37°C. Following incubation, 100 μl of diluted proteins + SARS-CoV-2 were transferred on Vero CCL-81 or E6 cell monolayer in 96-well plates with 80-90% confluency prepared a day prior to infection in duplicates and incubated at 37°C for 24 hours, and they were prepared for immunofluorescence staining.

### Immunofluorescence Staining

Vero CCL-81 or E6 cells were washed with 1X DPBS-T and fixed with 4% paraformaldehyde for 20 min at room temperature. Following three washes with 1X DPBS-T, the cells were permeabilized with 0.1% Triton X 100 in 1X PBS for 5 min and blocked with Protein Block (Abcam, USA) for 20 min at room temperature. Subsequently, the samples were incubated with rabbit anti-Spike (Sinobiological, #40589-T62) antibody for 90 min at 37°C. Following three further washes with 1X DPS-T, the cells were incubated with an appropriate secondary antibody (Alexa Fluor 488 goat anti-rabbit antibody, Thermo Scientific, USA) for 90 min at 37°C, and the nuclei were stained with Hoechst (Thermo Fisher Technology, #33342). The images were acquired with Leica DMi8 (Leica, Germany) live cell imaging microscope and were analyzed using Las X software.

### Spike binding assay

HEK293T cells were transfected with Spike-18aa truncated (Addgene plasmid # 149541), or engineered Spike-variants of concern (Alpha, Beta, Gamma, Delta) to express Spike protein in their cell surface. In parallel, HEK293T cells were transfected with sfGFP- or mRFP-fused versions of sACE2(WT), sACE2(v1) with or without SunTag and MoonTag system components. Spike-expressing HEK293T cells were seeded on 96-well plates and incubated with CM that were collected from cells transfected with sfGFP-sACE2 fusions, after 48h of transfection. For SunTag samples, 1 volume of CM from sACE2-5xGCN4 was mixed with 3 volumes of CM from sACE2-scFv_GCN4, and for MoonTag samples, 1 volume of CM from sACE2-12xgp41 was mixed with 4 volumes of CM from sACE2-nb_gp41 as the total volume is equal with CM from untagged sACE2s. For co-localization with SARS-CoV-2 Spike protein, cells were fixed and immunostained with anti-SARS-CoV-2 spike primary antibody, and Alexa Fluor-594 secondary antibody as explained in the SARS-CoV-2 neutralization assay. The images were acquired with Leica dmi8 live cell imaging microscope and were analyzed using Las X software.

### Animal studies

All experimental procedures with animals were approved by TUBITAK, Marmara Research Center, Genetic Engineering and Biotechnology Institute (TUBITAK MRC GEBI). All procedures in this study involving animals were reviewed and approved by the Institutional Biosafety Committee and Institutional Animal Care and Use Committee (HADYEK-16563500-111-58); all the experiments were complied with relevant ethical regulations. The experiments were conducted in Biosafety Level 3 (BSL3) and animal BSL3 (ABSL3) facilities at TUBITAK MRC GEBI. Jackson Laboratory in the United States provided the K18-hACE2 [B6.Cg-Tg(K18-hACE2)2Prlmn/J] transgenic mice used in this study. The TUBITAK MRC GEBI Experimental Animals unit is responsible for production of K18-hACE2 transgenic mice. All of the experiments were conducted in a biocontainment isocage, which is part of ABSL 3. 8-10 week old male K18-hACE2 transgenic mice were used in each group (n=8). There were two groups in this study: sACE2(v1)-MoonTag (20 mg/kg/day), and placebo (PBS).

The B.1.1.7 strain (alpha) of SARS-CoV-2 virus was employed in this study, which had 10^5^ TCID50 values. 50 μl of 10^5^ TCID50 SARS-CoV-2 virus was delivered intranasally (in) with under anesthesia for 3 days successively. Beginning from 4h after the 3rd day of infection, either sACE2(v1)-MoonTag (20 mg/kg/day), or placebo (PBS) were given to 8 mice intraperitoneally, for 4 consecutive days. Mice were evaluated for morbidity (body weight) and mortality on a daily basis after being infected with the virus. Mice showing >25 percent loss of their initial body weight were defined as reaching experimental endpoint and sacrificed. Gross pathologic examination was performed after the animals were sacrificed. Lungs were harvested for histopathologic evaluation. Half of the tissues were fixed in 10% neutral buffered formalin solution for pathology analyses. Briefly, fixed tissues were prepared in graded alcohol and xylene series, followed by paraffin embedding and sectioning. Sections were stained by Hematoxylin-eosin. The micrographs were obtained via use of a Zeiss Axio Lab A1 microscope (Zeiss, Germany) and zen blue software (Zeiss, Germany). The other half of the tissues were homogenized in 3mL of PBS using an ultrasonic homogenizer %70 amplitude for 90 seconds (BANDELIN HD2200.2) for viral isolation. Tissue homogenates were centrifuged at 17000 × g for 10 min and supernatants were collected to 15 ml falcon tubes.

### Viral RNA isolation and qRT-PCR

Viral RNA was extracted with the QIAamp Viral RNA Mini kit Cat: 52906 (QIAGEN) according to the protocols. The viral RNA quantification was performed using One Step PrimeScript III RT-qPCR Kit (Takara). All reactions were performed on a CFX96 Touch instrument with the following quantitative-PCR conditions: 52°C for 5 min, 95°C for 10 sec, followed by 44 cycles at 95°C for 5 sec and 55°C for 30 sec. The CDS primer sequences used for RT-qPCR were targeted against the Nucleocapsid (N) gene of SARS-CoV-2 with the following primers and probes: N1 Forward: 5’-GACCCCAAAATCAGCGAAAT-3’, N1 Reverse: 5’-TCTGGTTACTGCCAGTTGAATCTG-3’ N1 Probe: 5’-FAM-ACCCCGCATTACGTTTGGTGGACC-BHQ1-3 N2 Forward: 5’-TTACAAACATTGGCCGCAAA-3’ N2 Reverse: 5’-GCGCGACATTCCGAAGAA-3’ N2 Probe: 5’-FAM-ACAATTTGCCCCCAGCGCTTCAG-BHQ1-3.

### Statistical analysis

All data were analyzed with either GraphPad Prism, R studio or ImageJ softwares. Data were presented as mean +/- SD. An unpaired student’s t-test was used for comparison between two groups, while one way ANOVA was used for comparisons including multiple parameters. A two-sided p value < 0.05 was considered statistically significant. Details of each analysis are indicated in figure legends.

## Supplementary Materials

Fig. S1. Representative models for each fusion protein generated in this study

Fig. S2. Optimization of pseudovirus infection and ACE2-based neutralization

Fig. S3. Neutralization assay with SARS-CoV-2 Spike-bearing pseudovirus

Fig. S4. Neutralization assay with low dose SARS-CoV-2 isolate

Fig. S5. Spike-binding assay with GFP-fused sACE2 fusions

Fig. S6. Variant-Spike binding assay with sACE2 (WT)-MoonTag

Fig. S7. Representative histological images of lung sections harvested from each animal in the control or v1-MoonTag groups

Table S1. Details of vectors generated in this study.

Data file S1. Sequences of plasmids generated in this study.

Movie S1. Binding of v1-GFP-MoonTag to Spike-expressing HEK293T cells.

## Acknowledgments

We thank Dr. Tamer Onder for critical comments on the manuscript.

## Author contributions

Study design: AK, UA, DO and TBO; cloning: AK, UA and DO; pseudovirus production and neutralization: AK and CB; protein purification: AK, ES, NP, UA, CA and CB; SARS-CoV-2 isolation and neutralization: FC, OD, GE, TB, EN and BO; spike binding assay and IF staining: GNS and AK; *in vivo* experiments: HP and ISY; tissue preparation, staining and evaluation: GNS, GS and SK; supplied reagents and lines: NL, CA, MK, FC, and IS; data interpretation: AK, NL, MK, IS and TBO; graphics and figure design: NP, GNS, CB, and DO; initial manuscript draft: AK and TBO; approved final manuscript: all authors.

## Data and materials availability

All data are available in the main text or the supplementary materials. Genomic sequence information of clinical SARS-CoV-2 isolates used in this study is available with the accession number of GenBank: MT675956. Sequences of plasmids generated in this study can be found in Data File S1. All plasmids generated in this study are available to the scientific community by contacting AK and TBO, and will be available on the Addgene plasmid repository platform after publication.

