## Supplemental figures 1-7, Supplemental Table 1 for "Protein scaffold-based multimerization of soluble ACE2 efficiently blocks SARS-CoV-2 infection *in vitro* and *in vivo*"

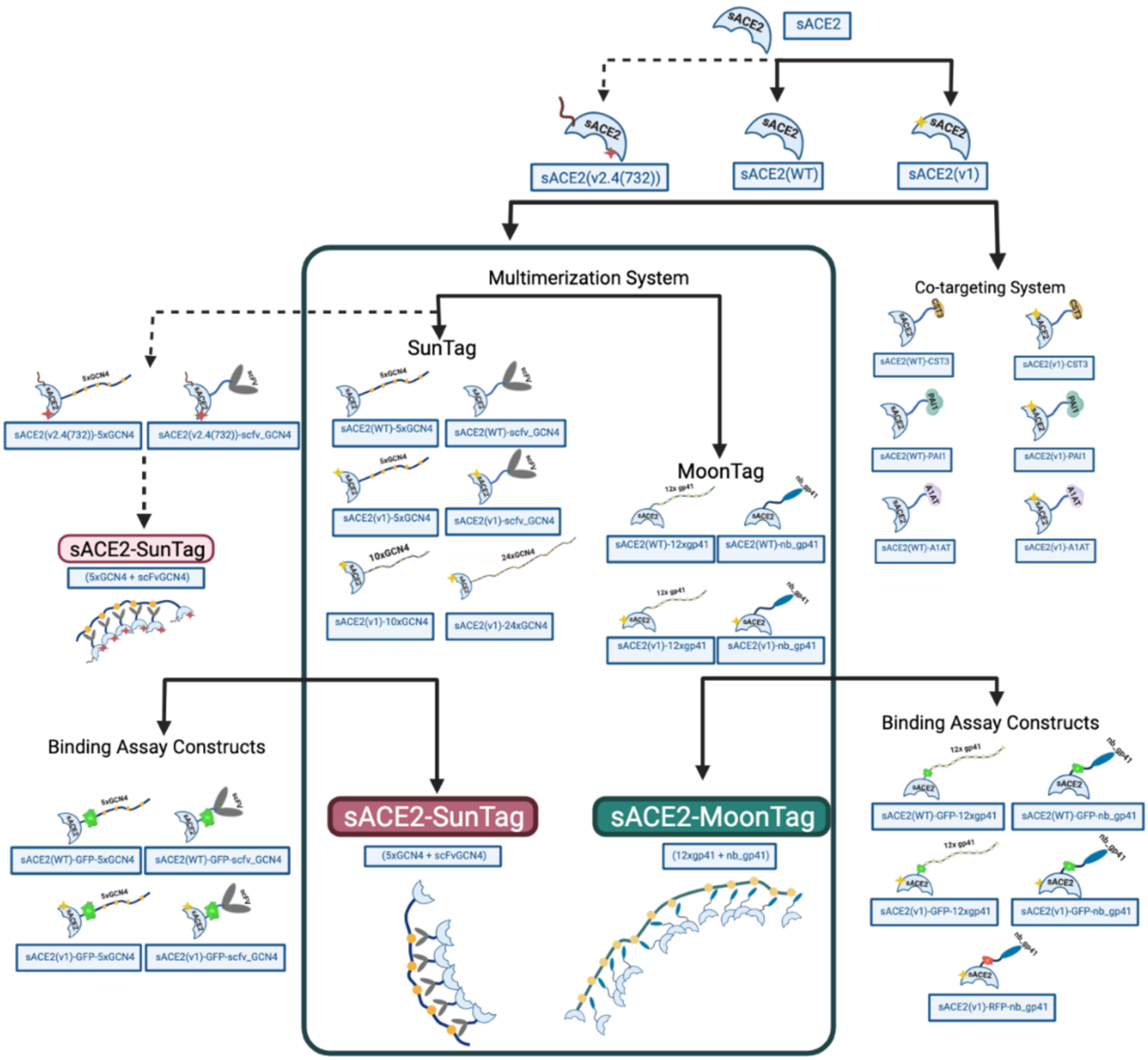


**Figure S1. Representative models for each fusion protein generated in this study.** *(Created with Biorender.com)*

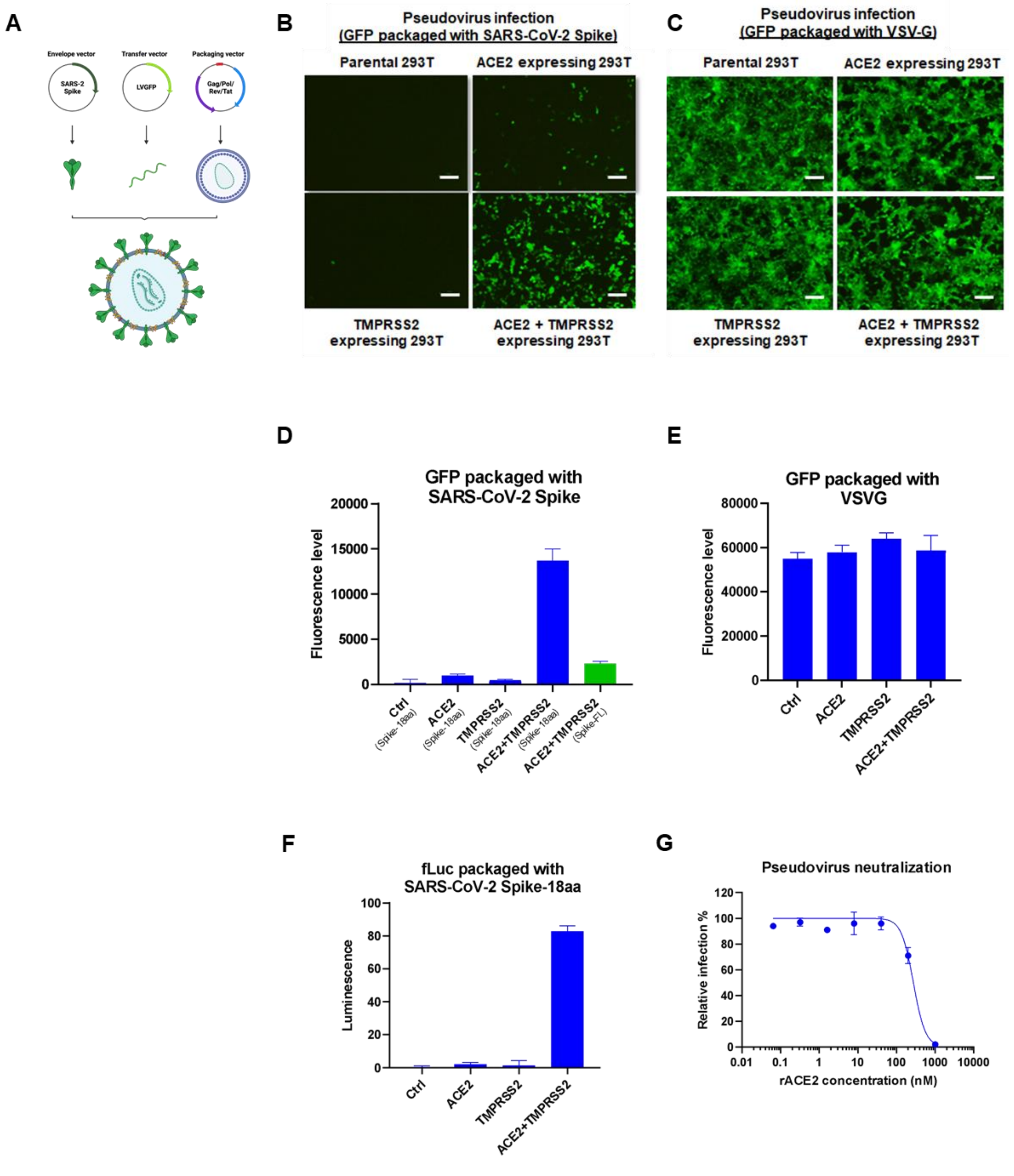


**Figure S2. Optimization of pseudovirus infection and ACE2-based neutralization. A)** Schematic representation of production of GFP-packaged pseudoviruses bearing SARS-CoV-2 Spike. *(Created with Biorender.com)* **B-C)** Microscopic images of infection of HEK293T cells expressing either ACE2 or TMPRSS2 individually, or ACE2+TMPRSS2 together, with GFP-packaged pseudoviruses bearing either SARS-CoV-2 Spike **(B)** or VSV-G **(C)** glycoproteins.

Scale bars = 100 µm. **D-E)** Quantification of infection with GFP-packaged pseudoviruses bearing SARS-CoV-2 Spike (18-aa deleted or full length) **(D)** or VSV-G **(E)** Parental HEK293T infection was used as control, and infection rates were normalized to fluorescence level of control. **F)** Quantification of infection with fLuc-packaged pseudoviruses bearing SARS-CoV-2 Spike. Parental HEK293T infection was used as control, and infection rates were normalized to luminescence level of control. **G)** Relative infection rate of ACE2 and TMPRSS2 expressing HEK293T cells with pseudoviruses bearing SARS-CoV-2 Spike (18-aa) in the presence of human recombinant ACE2 (rACE2).

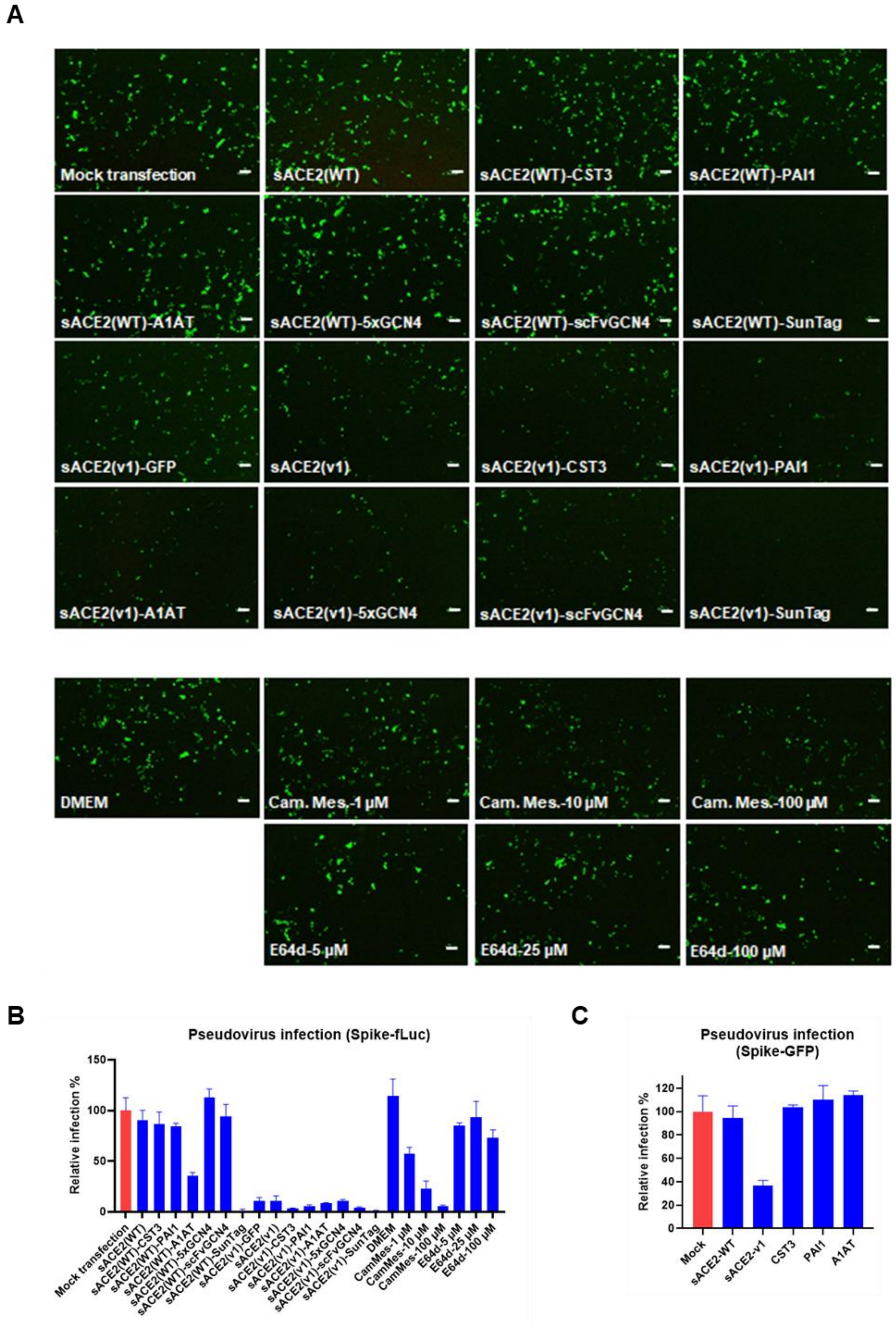


**Figure S3. Neutralization assay with SARS-CoV-2 Spike-bearing pseudovirus. A)** Microscopic images of HEK293T cells infected with GFP-packaged pseudoviruses in the presence of CM collected from sACE2-expressing or control cells, and chemicals camostat mesylate (CamMes) and E64d. Scale bar = 50 µm. **B)** Relative infection rate of HEK293T cells with fLuc-packaged pseudoviruses in the presence of CM collected from sACE2-expressing or mock transfected cells, and chemicals CamMes or E64d. CM from mock transfected cells were used as control, and infection rates were calculated as relative fluorescence to control wells. **C)** Relative infection of HEK293T cells with GFP-packaged pseudoviruses in the presence of CM collected from cells expressing sACE2, or several proteases (CST3, PAI1, A1AT) individually. CM from mock transfected cells were used as control, and infection rates were calculated as relative fluorescence to control wells.

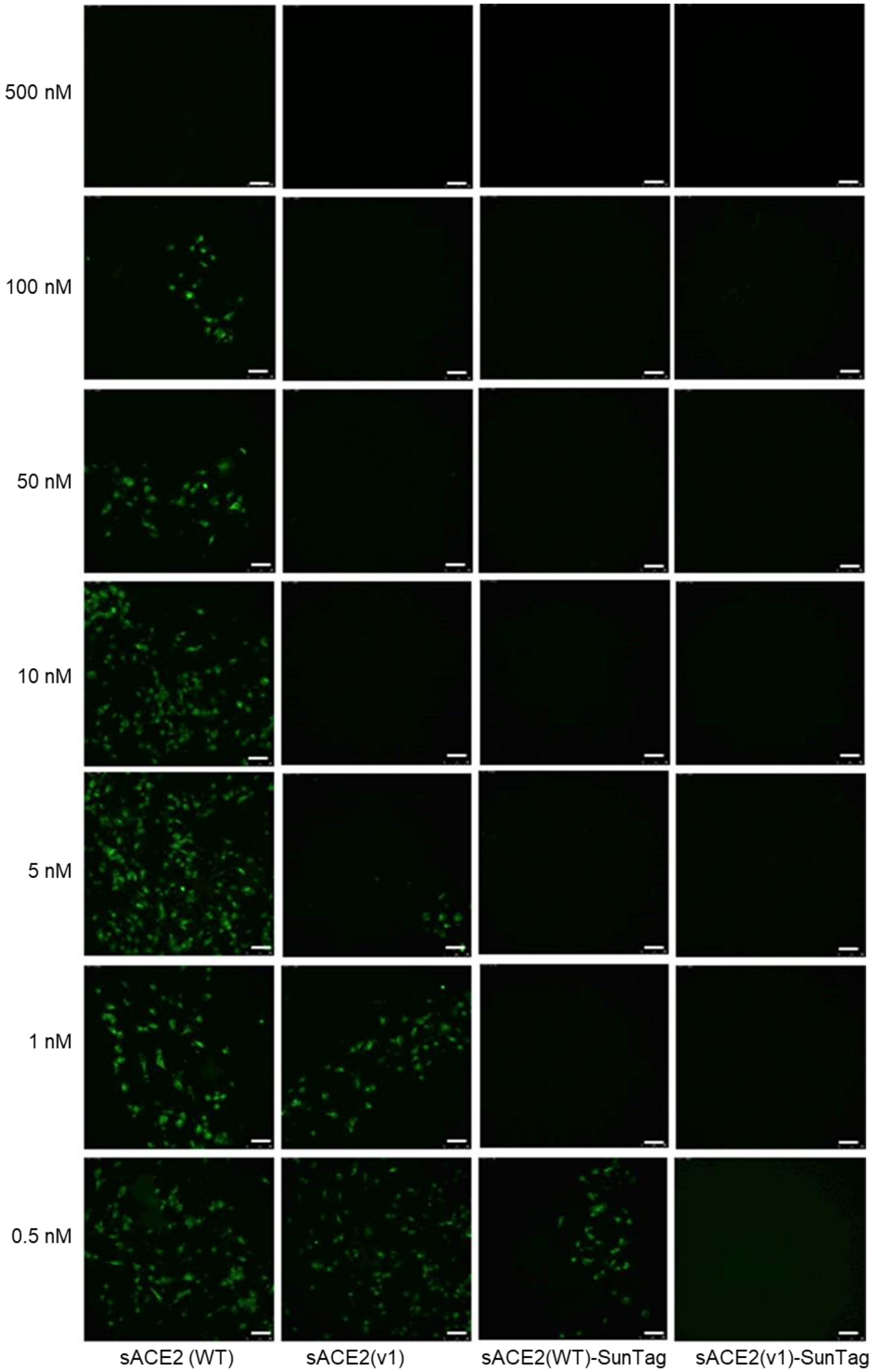


**Figure S4. Neutralization assay with low dose SARS-CoV-2 isolate.** Microscopic images of Vero-CCL81 cells infected with authentic SARS-CoV-2 ($10^3$ pfu/ml) upon incubation with different concentrations of purified sACE2 proteins for 1h. Cells were immunostained with anti-SARS-CoV-2 spike primary antibody (Green) after 24h of infection. Scale bar = 100 μm.

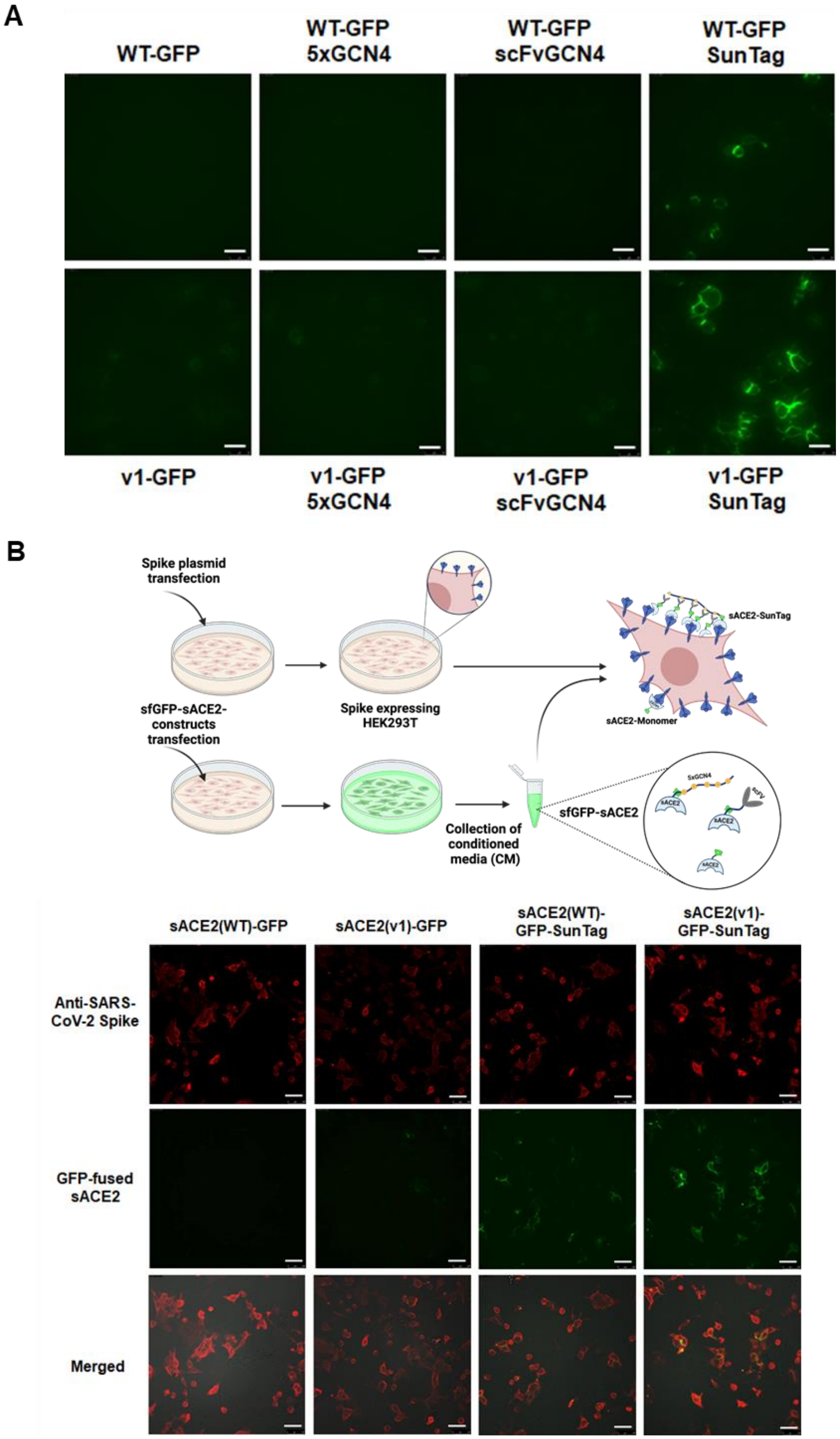


**Figure S5. Spike-binding assay with GFP-fused sACE2 fusions. A)** Microscopic images of Spike-expressing HEK293T cells upon incubation with CM that were collected from cells transfected with sfGFP-fused sACE2(WT) or sACE2(v1) with individual or combined SunTag components. Scale bars = 50 μm. **B)** Microscopic images of HEK293T cells expressing SARS-CoV-2 Spike upon incubation with CM collected from sfGFP fused sACE2(WT) or sACE2(v1) with or without SunTag system. Cells were immunostained with anti-SARS-CoV-2 antibody (red). Scale bars = 50 μm. *(Created with Biorender.com)*

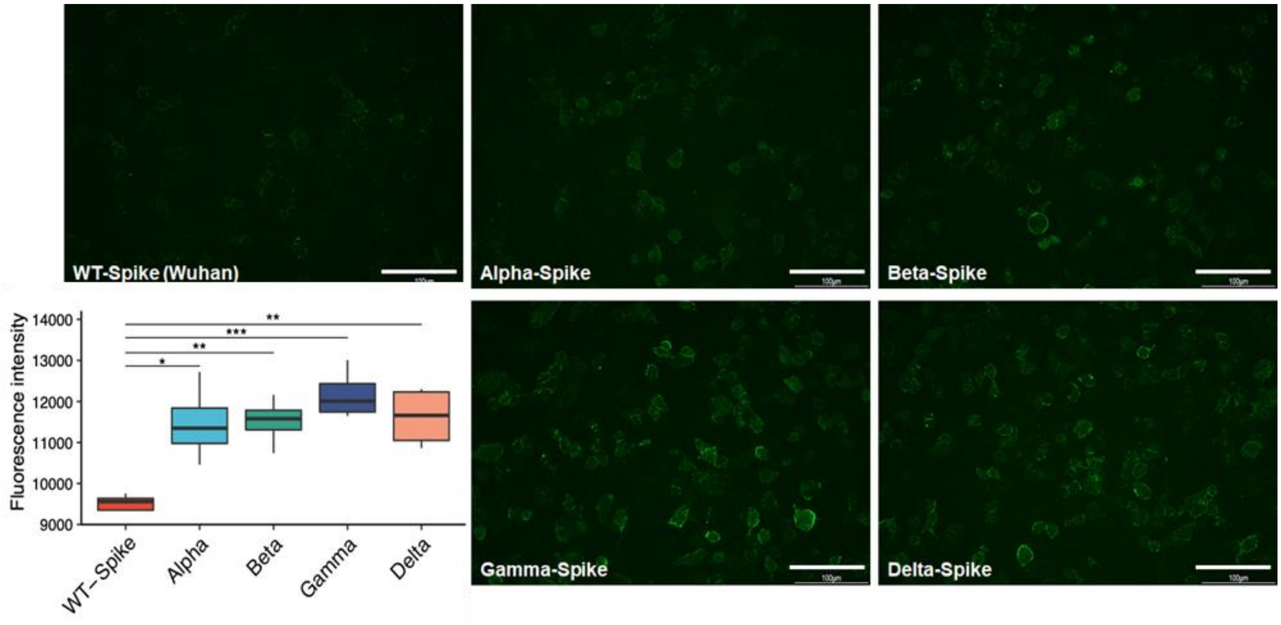


**Figure S6. Variant-Spike binding assay with sACE2 (WT)-MoonTag.** Microscopic images of wild type or VOC Spike-expressing HEK293T cells upon incubation with CM that were collected from cells transfected with sfGFP-fused version of sACE2(WT)-MoonTag. Scale bars = 100 µm.

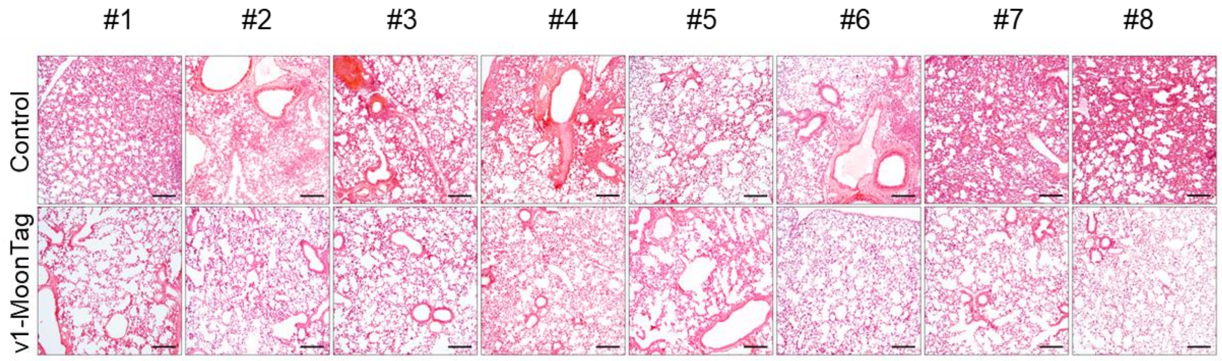


**Figure S7. Representative histological images of lung sections harvested from each animal in the control or v1-MoonTag groups.** Tissues obtained from v1-MoonTag groups have shown remarkable healing and restoration of tissue architecture in most of the animals. Scale bars=90 μm.

| Vector ID | Insert | Insert Size (bp) (Insert1-Linker-Insert2) | Linker | Purif. Tag (C-terminal) | Mutations |
|---|---|---|---|---|---|
| pcDNA3.1_sACE2(WT)-8h | sACE2(WT) | 1845 | | 8xHis | |
| pcDNA3.1_sACE2(WT)-CST3 | sACE2(WT)-CST3 | 1845+30+438 | GSGGSGSGGS | | |
| pcDNA3.1_sACE2(WT)-CST3-6h | sACE2(WT)-CST3 | 1845+30+438 | GSGGSGSGGS | 8xHis | |
| pcDNA3.1_sACE2(WT)-PAI1 | sACE2(WT)-PAI1 | 1845+30+1137 | GSGGSGSGGS | | |
| pcDNA3.1_sACE2(WT)-PAI1-6h | sACE2(WT)-PAI1 | 1845+30+1137 | GSGGSGSGGS | 6xHis | |
| pcDNA3.1_sACE2(WT)-A1AT | sACE2(WT)-A1AT | 1845+30+1182 | GSGGSGSGGS | | |
| pcDNA3.1_sACE2(WT)-A1AT-6h | sACE2(WT)-A1AT | 1845+30+1182 | GSGGSGSGGS | 6xHis | |
| pcDNA3.1_sACE2(WT)-5xGCN4 | sACE2(WT)-5xGCN4 | 1845+30+642 | GSNGPTDAAE | | |
| pcDNA3.1_sACE2(WT)-5xGCN4-8h | sACE2(WT)-5xGCN4 | 1845+30+642 | GSNGPTDAAE | 8xHis | |
| pcDNA3.1_sACE2(WT)-scFvGCN4 | sACE2(WT)-scFvGCN4 | 1845+30+741 | GSGGSGSGGS | | |
| pcDNA3.1_sACE2(WT)-scFvGCN4-8h | sACE2(WT)-scFvGCN4 | 1845+30+741 | GSGGSGSGGS | 8xHis | |
| pcDNA3.1_sACE2(WT)-sfGFP-5xGCN4 | sACE2(WT)-sfGFP-5xGCN4 | 1845+30+708+30+642 | GSGGSGSGGS/ GSNGPTDAAE | | |
| pcDNA3.1_sACE2(WT)-sfGFP-scFvGCN4 | sACE2(WT)-sfGFP-scFvGCN4 | 1845+30+708+30+741 | GSGGSGSGGS/ GSGGSGSGGS | | |
| pcDNA3.1_sACE2(WT)-12xgp41-8h | sACE2(WT)-12xgp41 | 1845+30+705 | GSGGSGSGSG | 8xHis | |
| pcDNA3.1_sACE2(WT)-nb_gp41-8h | sACE2(WT)-nb_gp41 | 1845+30+366 | GSGGSGSGGS | 8xHis | |
| pcDNA3.1_sACE2(WT)-sfGFP-12xgp41-8h | sACE2(WT)-sfGFP-12xgp41 | 1845+30+708+30+705 | GSGGSGSGGS/ GSGGSGSGSG | 8xHis | |
| pcDNA3.1_sACE2(WT)-sfGFP-nb_gp41-8h | sACE2(WT)-sfGFP-nb_gp41 | 1845+30+708+30+366 | GSGGSGSGGS/ GSGGSGSGGS | 8xHis | |
| pcDNA3.1_sACE2(v1)-8h | sACE2(v1) | 1845 | | 8xHis | H34A, T92Q, Q325P, A386L |
| pcDNA3.1_sACE2(v1)-CST3 | sACE2(v1)-CST3 | 1845+30+438 | GSGGSGSGGS | | H34A, T92Q, Q325P, A386L |
| pcDNA3.1_sACE2(v1)-CST3-6h | sACE2(v1)-CST3 | 1845+30+438 | GSGGSGSGGS | 6xHis | H34A, T92Q, Q325P, A386L |
| pcDNA3.1_sACE2(v1)-PAI1 | sACE2(v1)-PAI1 | 1845+30+1137 | GSGGSGSGGS | | H34A, T92Q, Q325P, A386L |
| pcDNA3.1_sACE2(v1)-PAI1-6h | sACE2(v1)-PAI1 | 1845+30+1137 | GSGGSGSGGS | 6xHis | H34A, T92Q, Q325P, A386L |
| pcDNA3.1_sACE2(v1)-A1AT | sACE2(v1)-A1AT | 1845+30+1182 | GSGGSGSGGS | | H34A, T92Q, Q325P, A386L |
| pcDNA3.1_sACE2(v1)-A1AT-6h | sACE2(v1)-A1AT | 1845+30+1182 | GSGGSGSGGS | 6xHis | H34A, T92Q, Q325P, A386L |
| pcDNA3.1_sACE2(v1)-5xGCN4 | sACE2(v1)-5xGCN4 | 1845+30+642 | GSNGPTDAAE | | H34A, T92Q, Q325P, A386L |
| pcDNA3.1_sACE2(v1)-5xGCN4-8h | sACE2(v1)-5xGCN4 | 1845+30+642 | GSNGPTDAAE | 8xHis | H34A, T92Q, Q325P, A386L |
| pcDNA3.1_sACE2(v1)-scFvGCN4 | sACE2(v1)-scFvGCN4 | 1845+30+741 | GSGGSGSGGS | | H34A, T92Q, Q325P, A386L |
| pcDNA3.1_sACE2(v1)-scFvGCN4-8h | sACE2(v1)-scFvGCN4 | 1845+30+741 | GSGGSGSGGS | 8xHis | H34A, T92Q, Q325P, A386L |
| pcDNA3.1_sACE2(v1)-10xGCN4 | sACE2(v1)-10xGCN4 | 1845+30+1398 | GSGGSGSGGS | | H34A, T92Q, Q325P, A386L |
| pcDNA3.1_sACE2(v1)-24xGCN4 | sACE2(v1)-24xGCN4 | 1845+30+1782 | GSGGSGSGGS | | H34A, T92Q, Q325P, A386L |
| pcDNA3.1_sACE2(v1)-sfGFP-5xGCN4 | sACE2(v1)-5xGCN4 | 1845+30+708+30+642 | GSGGSGSGGS/ GSNGPTDAAE | | H34A, T92Q, Q325P, A386L |
| pcDNA3.1_sACE2(v1)-sfGFP-scFvGCN4 | sACE2(v1)-scFvGCN4 | 1845+30+708+30+741 | GSGGSGSGGS/ GSGGSGSGGS | | H34A, T92Q, Q325P, A386L |
| pcDNA3.1_sACE2(v1)-12xgp41-8h | sACE2(v1)-12xgp41 | 1845+30+705 | GSGGSGSGSG | 8xHis | H34A, T92Q, Q325P, A386L |
| pcDNA3.1_sACE2(v1)-nb_gp41-8h | sACE2(v1)-nb_gp41 | 1845+30+366 | GSGGSGSGGS | 8xHis | H34A, T92Q, Q325P, A386L |
| pcDNA3.1_sACE2(v1)-sfGFP-12xgp41-8h | sACE2(v1)-sfGFP-12xgp41 | 1845+30+708+30+705 | GSGGSGSGGS/ GSGGSGSGSG | 8xHis | H34A, T92Q, Q325P, A386L |
| pcDNA3.1_sACE2(v1)-sfGFP-nb_gp41-8h | sACE2(v1)-sfGFP-nb_gp41 | 1845+30+708+30+366 | GSGGSGSGGS/ GSGGSGSGGS | 8xHis | H34A, T92Q, Q325P, A386L |
| pcDNA3.1_sACE2(v1)-mRFP-nb_gp41-8h | sACE2(v1)-mRFP-nb_gp41 | 1845+30+672+30+366 | GSGGSGSGGS/ GSGGSGSGGS | 8xHis | H34A, T92Q, Q325P, A386L |
| pcDNA3.1_sACE2v2.4(732)-5xGCN4-8h | sACE2v2.4(732)-5xGCN4 | 2193+30+642 | GSNGPTDAAE | 8xHis | T27Y, L79T, N330Y |
| pcDNA3.1_sACE2v2.4(732)-scFvGCN4-8h | sACE2v2.4(732)-scFvGCN4 | 2193+30+741 | GSGGSGSGGS | 8xHis | T27Y, L79T, N330Y |
| SARS-CoV-2 Spike-Alpha (C-term. 18-aa trunc.) | SARS-CoV-2 Spike | 3759 | | | H69V70 del, N501Y, D614G, P681H |
| SARS-CoV-2 Spike-Beta (C-term. 18-aa trunc.) | SARS-CoV-2 Spike | 3765 | | | K417N, E484K, N501Y, D614G |
| SARS-CoV-2 Spike-Gamma (C-term. 18-aa trunc.) | SARS-CoV-2 Spike | 3765 | | | K417T, E484K, N501Y, D614G |
| SARS-CoV-2 Spike-Delta (C-term. 18-aa trunc.) | SARS-CoV-2 Spike | 3765 | | | L452R, T478K, D614G, P681R |

**Table S1. Details of vectors generated in this study**
